# Senescent Activated Naive B Cells Promote Anti-Citrullinated Antigen T Cell Responses and the Transition to Clinical Rheumatoid Arthritis

**DOI:** 10.1101/2025.10.14.682430

**Authors:** Xiaohao Wu, Mengrui Zhang, Jae-Seung Moon, Eun Kyung Song, Joshua C. Abrams, Laura S. van Dam, Orr Sharpe, Marie Feser, Laurie Moss, Peggy P. Ho, Melanie H. Smith, Laura T. Donlin, Jane H. Buckner, Eddie A. James, Gary S. Firestein, Yuko Okamoto, Tobiaz V. Lanz, Eric Meffre, V. Michael Holers, Kevin D. Deane, William H. Robinson

## Abstract

Rheumatoid arthritis (RA) is a chronic autoimmune disease marked by joint and systemic inflammation. Anti-citrullinated protein antibodies (ACPAs) define an at-risk stage that precedes clinically apparent inflammatory arthritis (clinical RA) onset, yet the molecular mechanisms driving progression remain poorly understood. Here, we applied single-cell multi-omics to profile B cells longitudinally collected from ACPA⁺ individuals who either convert to clinical RA (Converters) or do not (Nonconverters). We identified a striking expansion of CXCR5⁺CD69⁺ activated naive B cells (aNAVs) uniquely in Converters prior to clinical RA. These aNAVs exhibited a pro-inflammatory, senescent transcriptional program and persist through to clinical RA. In Converters, aNAVs expressed polyreactive, autoreactive IgM with distinctive V–J gene rearrangements that dominate the BCR repertoire. Furthermore, in Converters most IgM⁺ aNAVs were developmentally arrested in the peripheral blood, while a subset undergoes class switching and follows divergent somatic hypermutation trajectories. Mechanistically, aNAVs infiltrated RA synovium and served as potent antigen presenting cells to activate both anti-citrullinated antigen CD4⁺ and CD8⁺ T cells in an HLA-dependent manner. Chronic exposure to citrullinated antigens and CpG synergistically drove aNAV activation and senescence. These findings establish a mechanistic link between naive B cell senescence and clinical RA development in ACPA^+^ individuals, providing a rationale for therapeutically targeting aNAV B cells for the prevention of RA.

**Graphic abstract:** 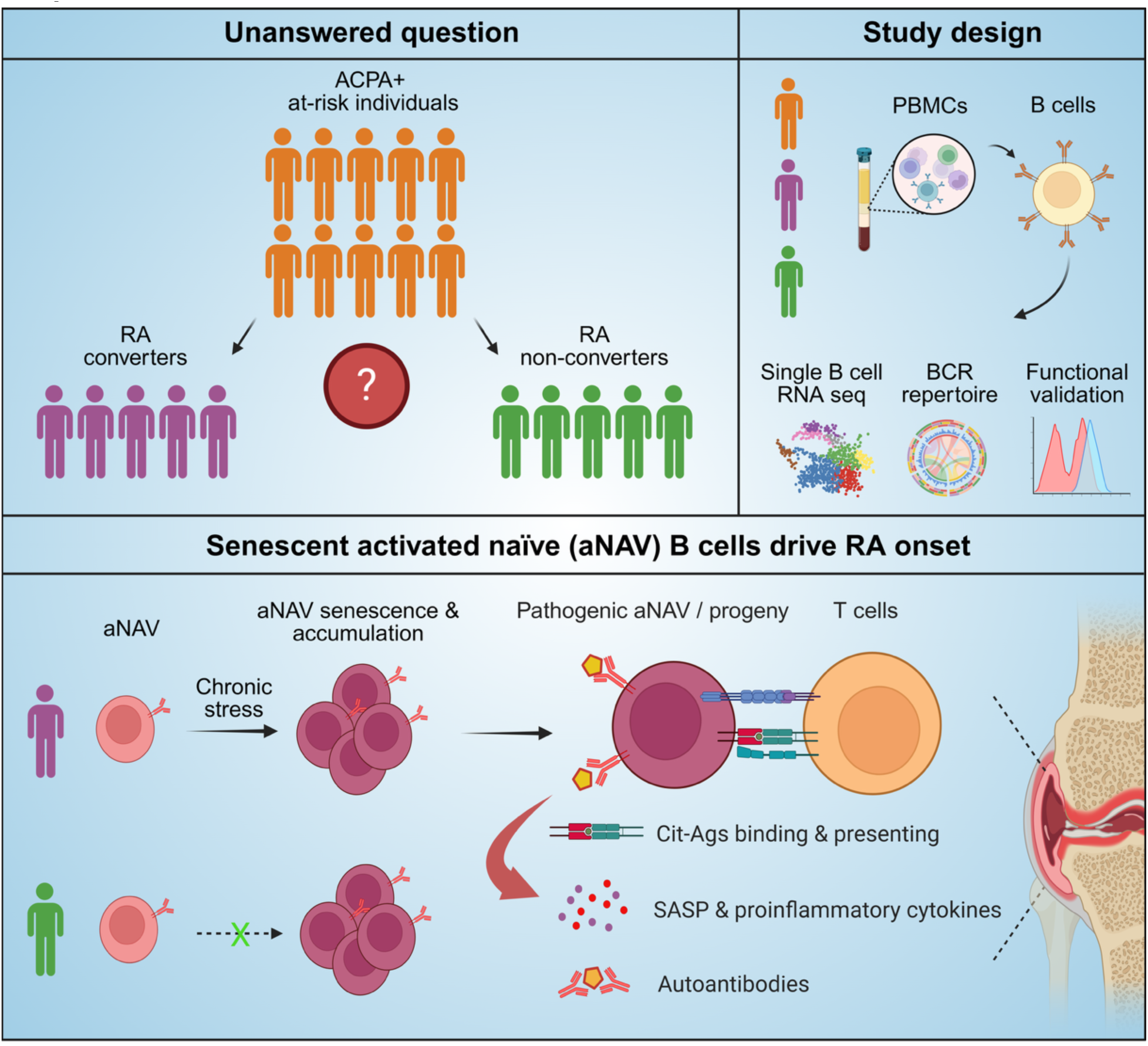

## Introduction

Rheumatoid arthritis (RA) is an chronic autoimmune disease marked by persistent synovial inflammation, progressive joint destruction, and systemic comorbidities [1]. Affecting ∼1% of the global population, RA imposes a significant burden through chronic pain, disability, reduced quality of life, and increased healthcare costs [2]. Diagnosis is based on clinical features, serological detection of autoantibodies, such as rheumatoid factor (RF) and anti-citrullinated protein antibodies (ACPAs), and imaging to assess joint involvement [3]. Current therapies, including disease-modifying antirheumatic drugs (DMARDs), cytokine- or immune cell–targeting biologics, and Janus kinase inhibitors, can control disease activity in many patients [4]. However, early diagnosis and prevention remain major unmet needs.

The well-established use of B cell directed therapy (e.g. rituximab) to treat RA, as well as recent success of anti-CD19 CAR T cell therapy in clinical trials for autoimmune diseases highlights the central role of B cells in autoimmunity, including RA [5–9]. Beyond their role as autoantibody-producing cells, B cells also serve as potent antigen-presenting cells, critically shaping T cell responses [10]. Although autoreactive B cells naturally arise in healthy individuals, they are typically eliminated by central and peripheral tolerance mechanisms [11, 12]. In RA, however, these tolerance checkpoints fail, allowing the survival and expansion of autoreactive B cells [13–15]. In this context, B cell–derived autoantibodies not only serve as diagnostic biomarkers but also functionally contribute to disease pathogenesis—either by promoting inflammation or, in some cases, by modulating immune responses to mitigate tissue damage [16–19].

ACPAs recognize citrullinated proteins and peptides generated by peptidylarginine deiminase (PAD)–mediated conversion of arginine to citrulline [20]. While citrullination occurs physiologically, aberrant or excessive citrullination, and autoimmune responses directed to citrullinated proteins, is implicated in RA pathogenesis [21]. ACPAs exhibit high specificity for RA (∼95–98%) compared to rheumatoid factor (RF; ∼80–90%). Importantly, it is now well-established that ACPA elevations in the blood can precede the development of clinically-apparent inflammatory arthritis and a diagnosis of RA (clinical RA) on average by 3–5 years [22–25]. The period of ACPA elevations without clinical RA can be termed an ‘at-risk’ stage of disease, and there are increasing efforts to target this stage with preventive interventions [26]. Recent studies have demonstrated that, during the at-risk stage, ACPA-positive individuals exhibit distinct epigenetic landscapes and immunophenotypic features which correlate with subsequent development of clinical RA [27–33], suggesting that RA may be a predictable and potentially preventable disease. Nevertheless, the cellular and molecular mechanisms underlying the transition from the ACPA-positive at-risk phase to clinical RA remain poorly understood.

In this study, we sought to further define the cellular and molecular mechanisms underlying the progression from preclinical to clinical RA by applying multi-modal single-cell sequencing to comprehensively profile circulating B cells from before and after the development of clinical RA. We analyzed longitudinally collected samples from prior to and after the onset of clinical RA from ACPA⁺ individuals who converted to clinical RA (Converters), as well as longitudinal samples from ACPA⁺ individuals who did not develop RA (Nonconverters), and ACPA⁻ healthy controls (Controls). The inclusion of ACPA⁺ non-converters as a comparator was important, enabling us to distinguish features specifically associated with development of clinical RA rather than ACPA positivity alone. Our integrative analyses revealed profound preclinical alterations in the B cell compartment, including shifts in cellular composition, transcriptional reprogramming, and remodeling of the B cell receptor (BCR) repertoire. These changes were marked by the expansion and pathogenic activation of a CXCR5⁺CD69⁺ naive (aNAV) B cell population prior to clinical RA. Together, our data suggest that aNAV B cells play a key role in the development of clinical RA by skewing the BCR repertoire toward autoreactivity and promoting anti-citrullinated antigen T cell responses.

## Results

### Comprehensive single-B-cell profiling of RA clinical conversion

In this study, we analyzed a sex- and age-matched cohort from the Targeting Immune Responses for Prevention of Rheumatoid Arthritis (TIP-RA) project. Participants were stratified into three groups based on ACPA status and subsequent clinical RA development, with longitudinal sampling from a single pre-clinical RA time point, and a clinical RA time point (Fig. 1a; Supplementary Table 1). The ACPA⁺ Converters included five individuals who were ACPA-positive at the first time point (i.e. pre-clinical RA) and subsequently developed clinical RA by the second. The ACPA⁺ Nonconverters included five individuals who were ACPA-positive at the first time point and had not developed clinical RA by their second time point. To avoid inadvertently including individuals who may have progressed to clinical RA shortly thereafter, we selected only those who remained RA-free for at least one year following their second sample. The ACPA⁻ Controls included three individuals who were negative for both ACPA and clinical RA at both time points. All participants were female to minimize sex effects. To minimize variability between participants in the time span of the temporal development of clinical RA, we selected a baseline sample for Converters that was approximately one-year prior to the onset of clinical RA, and a sample from the visit at which clinical RA was first identified. The mean interval between the two sampling timepoints was 470.6 ± 123.44 days for Converters, 340.8 ± 20.52 days for Nonconverters, and 337 ± 15.87 days for Controls. At baseline, anti-CCP3 (anti–cyclic citrullinated peptide antibodies) (Inova Diagnostics, Inc., San Diego, California USA) and C-reactive protein (CRP) levels were comparable between Converters and Nonconverters, and neither group had inflammatory arthritis. For this study, the outcome of clinical RA in Converters was defined as the presence of >=1 swollen joint on physical examination that also met the 2010 American College of Rheumatology/European Alliance of Associations for Rheumatology Classification Criteria for RA [34]. Peripheral blood was collected at each time point and PBMCs were isolated. B cells were enriched by negative selection and profiled using integrated 5’ single-cell RNA-seq (scRNA-seq), VDJ-seq, and CITE-seq. After stringent quality control, 165,247 B cells were retained for downstream analysis (Fig. 1b).

**Figure 1.**
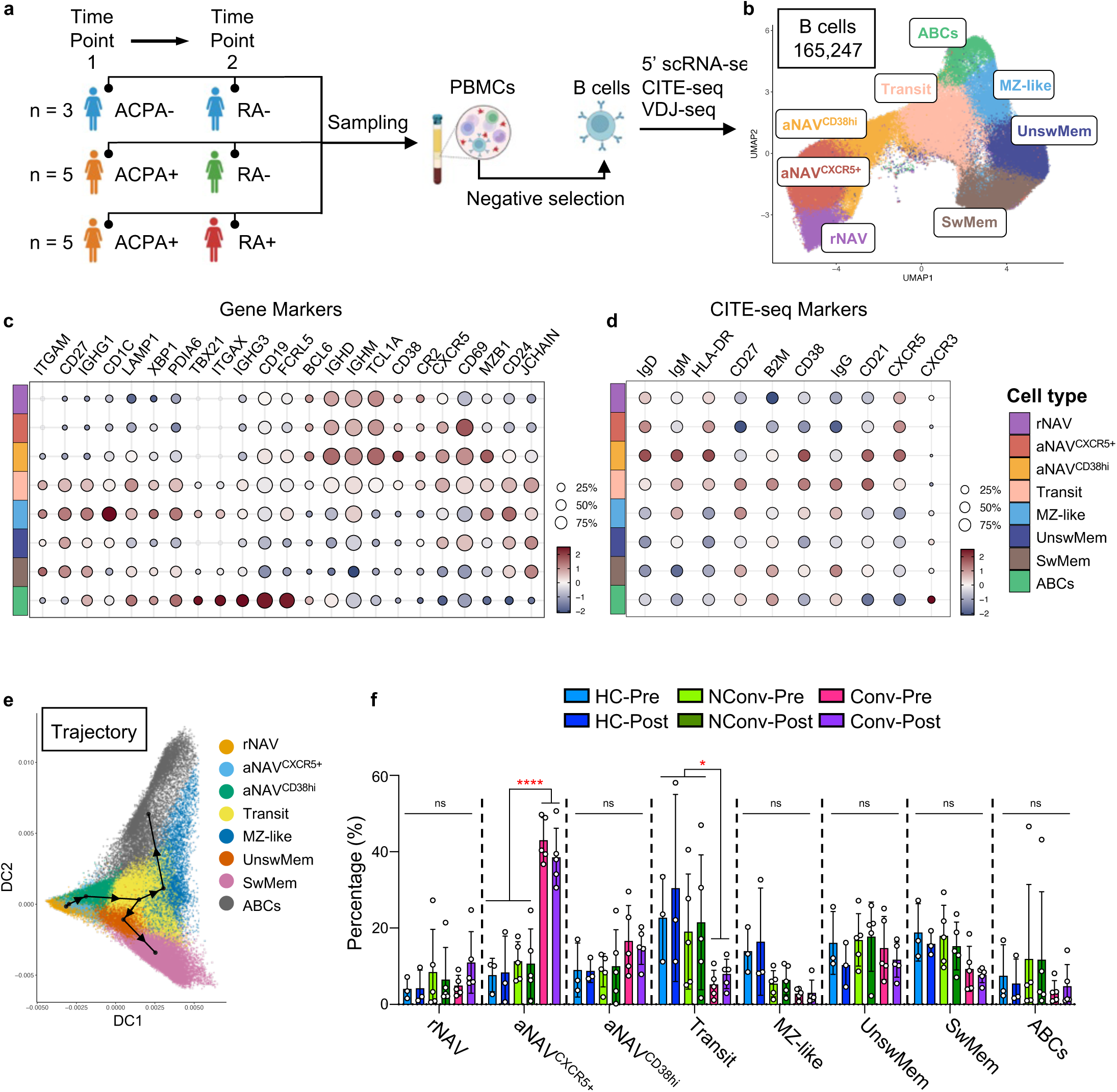
Single-B-cell profiling reveals a striking preclinical expansion of aNAV^CXCR5⁺^ cells in ACPA⁺ Converters. (**a**) Schematic of the study design. PBMCs were collected from ACPA⁺ Converters (n = 5), ACPA⁺ Nonconverters (n = 5), and ACPA-Controls (n = 3) at two time points. B cells were enriched by negative selection and profiled using integrated 5’ scRNA-seq, CITE-seq, and VDJ-seq. (**b**) UMAP projection of B cell subsets, including resting naive (rNAV), early-phase activated naive (aNAV^CXCR5⁺^), later-phase activated naive (aNAV^CD38hi^), transitioning B cells (Transit), marginal zone– like (MZ-like), unswitched memory (UnswMem), switched memory (SwMem), and atypical B cells (ABCs). A total of 165,247 high-quality B cells were retained after quality control and filtering. (**c**) Transcriptomic marker expression profiles across subsets. (**d**) CITE-seq quantification of surface protein expression. (**e**) Pseudotime trajectory analysis showing continuous differentiation from rNAV through aNAV, with divergence at the Transit cluster toward either MZ-like/ABC or memory B cell lineages. (**f**) Proportional analysis demonstrating a marked expansion of the aNAV^CXCR5⁺^ compartment, accompanied by a reduction in the Transit compartment, in Converters at both pre-clinical and post-onset stages. *P < 0.05, ****P < 0.0001 by two-way ANOVA (f); ns, not significant. HC: ACPA^-^ Controls; NConv: ACPA^+^ Nonconverters; Conv: ACPA^+^ Converters; Pre: First time point (pre-clinical); Post: Second time point (post-onset).

UMAP projection revealed distinct B-cell populations, including resting naive (rNAV), activated naive (aNAV), transitioning (Transit), marginal zone–like (MZ-like), unswitched memory (UnswMem), switched memory (SwMem), and atypical B cells (ABCs) (Fig. 1c). Marker gene expression delineated clear molecular signatures: naive cells were CD19⁺ CD27⁻ IGHM⁺ IGHD⁺ TCL1A⁺, whereas memory-like subsets expressed CD27 with variable IGHM and reduced IGHD. ABCs were CD19⁺ CD27⁻ ITGAX (CD11c)⁺ TBX21 (T-bet)⁺ FCRL5⁺ CD21^low^, and MZ-like cells expressed high CD1C and MZB1. CITE-seq measurements confirmed these surface phenotypes, supporting the transcriptomic annotations (Fig. 1d). Pseudotime analysis resolved a continuous differentiation trajectory originating from rNAV, passing through an aNAV intermediate, and bifurcating toward MZ-like/ABC versus UnswMem/SwMem lineages (Fig. 1e).

### Preclinical expansion of aNAV^CXCR5⁺^ cells in ACPA⁺ Converters

Proportional analysis showed a striking pre-clinical RA expansion of aNAV cells (CD19⁺ CD27⁻ IGHM⁺ IGHD⁺ CXCR5⁺ CD69⁺ CD38⁻) in individuals who progressed to clinical RA, relative to Nonconverters and Controls (Fig. 1f). This expansion persisted in Converter samples after clinical RA onset. In Converters, aNAVs constituted 43.01 ± 5.91% of B cells at the pre-clinical RA stage and 38.55 ± 7.60% post-onset. In Nonconverters, the corresponding proportions were 11.34 ± 5.10% (pre) and 10.68 ± 9.09% (post) (p < 0.001 for both comparisons versus Converters). aNAV^CXCR5⁺^ frequencies did not differ between Nonconverters and Controls. The expansion of aNAV in Converters was accompanied by a significant reduction in transitioning B cells at both time points (p < 0.05 versus Nonconverters and Controls), whereas other subsets were unchanged.

### Pathogenic activation of aNAV^CXCR5⁺^ cells in ACPA⁺ Converters

To characterize the naive compartment, naive B cells were resolved into three transcriptionally distinct clusters: one rNAV and two sequential aNAV states, distinguished by activation and BCR-signaling programs. rNAVs expressed the lowest levels of activation and BCR signaling genes; both aNAV clusters upregulated canonical activation markers (CD69, JUN, JUND, CD83, FOS, FOSB, JUNB, NFKB1, NFKB2) (Fig. 2a). CD38 distinguished early aNAV^CXCR5⁺^ from later aNAV^CD38hi^, the latter showing higher LYN and SYK, consistent with progressive activation.

**Figure 2.**
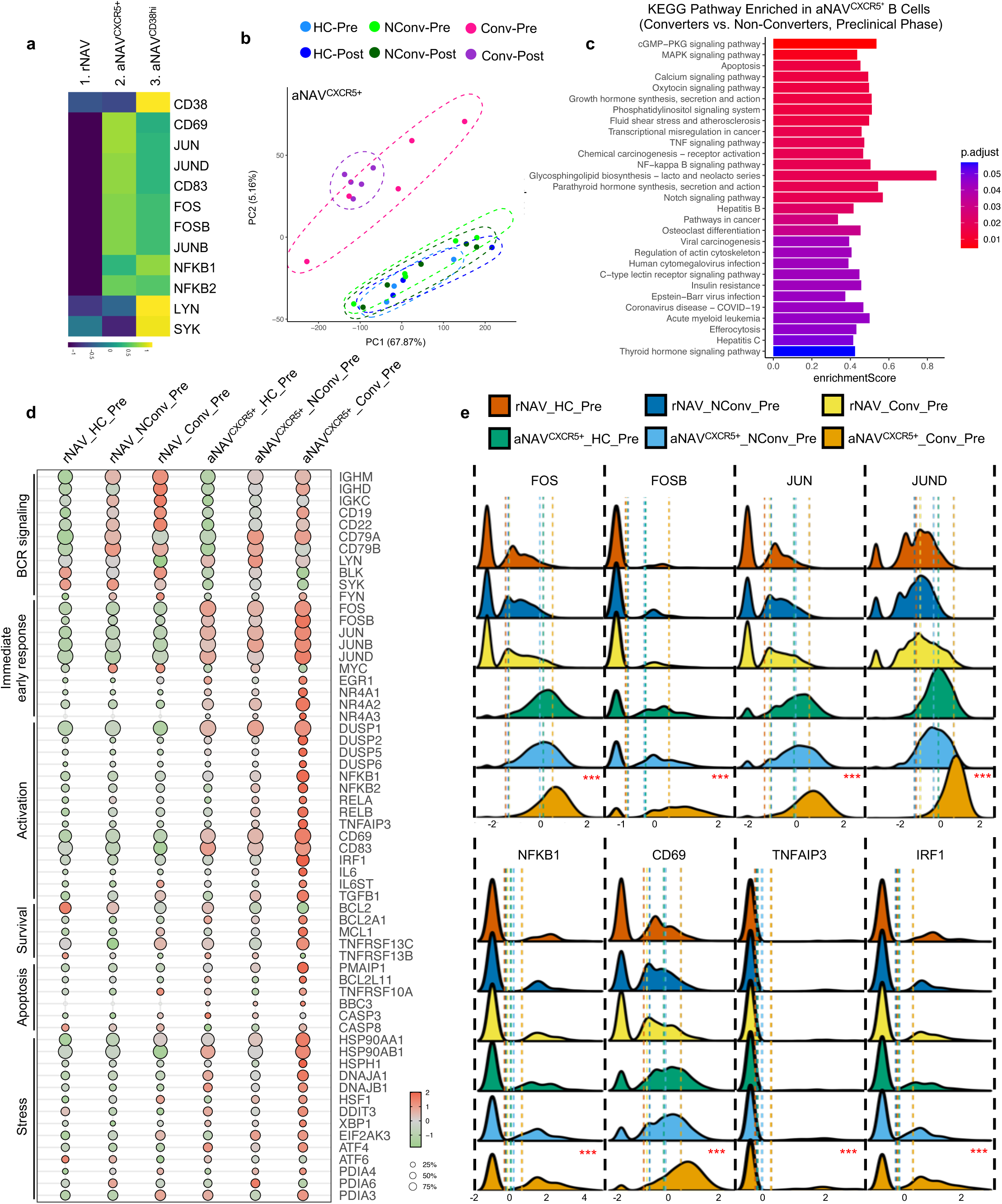
Converters’ aNAV^CXCR5⁺^ cells display proinflammatory, stressed, and anergy-like transcriptional profiles. (**a**) Expression of activation markers across three naive B cell clusters, showing elevated activation marker expression (e.g., CD69, JUN, JUND) in both activated naive clusters. CD38 distinguishes the early (aNAV^CXCR5⁺^) from the later (aNAV^CD38hi^) activation phase. (**b**) PCA analysis showing that the transcriptional profiles of aNAV^CXCR5⁺^ cells are distinct in Converters compared with Nonconverters and Controls. Each dot represents an individual patient sample. (**c**) KEGG pathway enrichment analysis of aNAV^CXCR5⁺^ cells comparing Converters and Nonconverters at the pre-clinical RA phase. The x-axis indicates enrichment scores; color denotes significance (adjusted P value). (**d**) Dot plots showing expression of genes grouped by functional categories, including BCR signaling, immediate early response, activation, survival, apoptosis, and cellular stress. (**e**) Ridge plots illustrating the distribution of representative activation genes (FOS, FOSB, JUN, JUND, NFKB1, CD69, TNFAIP3, IRF1) in rNAV and aNAV^CXCR5⁺^ cells across the three groups. Dashed lines indicate mean expression levels. ***P < 0.001 by Kruskal–Wallis test followed by Dunn’s post hoc test for comparison of aNAV^CXCR5⁺^ cells between Converters and all other groups (e). rNAV: resting naive; aNAV: activated naive; HC: ACPA^-^ Controls; NConv: ACPA^+^ Nonconverters; Conv: ACPA^+^ Converters; Pre: First time point (pre-clinical); Post: Second time point (post-onset).

Focusing on the expanded aNAV^CXCR5⁺^ population, PCA revealed transcriptional profiles in Converters that were distinct in samples from both prior to and after the onset of clinical RA when compared to longitudinal samples from Nonconverters and Controls; furthermore, the profiles from the Nonconverters and Controls largely overlapped (Fig. 2b). KEGG pathway analysis of pre-clinical RA aNAV^CXCR5⁺^ cells from Converters revealed significant enrichment of inflammatory signaling pathways, including TNF, MAPK, and NF- κB, as well as apoptosis and several virus-related pathways (e.g., hepatitis B, HCMV, EBV), compared to aNAV^CXCR5⁺^ cells from Nonconverters (Fig. 2c).

Gene-set analysis highlighted marked dysregulation emerging at the aNAV^CXCR5⁺^ stage in Converters (Fig. 2d). At the pre-clinical RA stage, Converters’ aNAV^CXCR5⁺^ cells upregulated upstream BCR components (IGHM, IGHD, CD19, CD22) but downregulated downstream mediators (LYN, BLK, SYK), suggestive of an anergy-like state under chronic low-level antigen engagement [35, 36]. Immediate-early genes (FOS, FOSB, JUN, JUNB, JUND, EGR1, NR4A1, NR4A2, NR4A3) were strongly induced, alongside MAPK/NF-κB activation (CD69, CD83, NFKB1, RELA, RELB) and cytokines (IL6, TGFB1). Survival programs shifted toward inflammation-adapted states (BCL2A1, MCL1, TNFRSF13C) with reduced BCL2. Stress and apoptotic mediators (PMAIP1, BCL2L11, HSP90AA1/AB1, DNAJA1/JB1, ATF4, XBP1, PDIA4) were elevated, indicating a highly stressed, checkpoint-resistant phenotype. Ridge plot analyses confirmed distinct distributions of activation/stress markers (e.g., FOS, FOSB, JUN, JUND, CD69) between aNAV^CXCR5⁺^ and rNAV subsets (Fig. 2e). Notably, Converters’ aNAV^CXCR5⁺^ cells expressed significantly higher FOS, FOSB, JUN, JUND, NFKB1, CD69, TNFAIP3, and IRF1 than their counterparts in both comparison groups (P < 0.001 by Kruskal–Wallis test followed by Dunn’s post hoc test), indicating a pathogenic autoreactive-like program before onset of clinical RA.

### Converters’ aNAV^CXCR5⁺^ exhibit senescence and developmental arrest

Gene Ontology analysis comparing pre-clinical RA aNAV^CXCR5⁺^ cells from Converters revealed significant enrichment of stress- and response-related pathways compared to Nonconverters, including the “Response to Mechanical Stimulus” pathway and the “Integrated Stress Response Signaling” pathway. In contrast, pathways associated with energy metabolism, such as aerobic respiration, oxidative phosphorylation, mitochondrial ATP synthesis, and cellular respiration, were consistently downregulated in aNAV^CXCR5⁺^ cells from Converters, indicative of a senescent metabolic phenotype. (Fig. 3a). In line with this, Converters’ aNAV^CXCR5⁺^ cells showed upregulation of stress and senescence-associated genes (Fig. 3b), including DNA damage and stress response mediators (HUS1, HRAS, RAF1), the master regulator CDKN1A (p21), SMAD3, cell-cycle repressors (RBL1, RBL2, E2F4, E2F5), SASP factors (TGFB1, IL6), and transcription factors KLF2, KLF4, and KLF6. Notably, these transcriptional alterations were already prominent at the pre-clinical RA stage in Converters. In contrast, Nonconverters’ aNAV^CXCR5⁺^ cells showed only modest upregulation of select genes, such as TP53, GADD45G, RBL2, and E2F5, at the second timepoint, suggesting a limited or transient activation of the senescence program.

**Figure 3.**
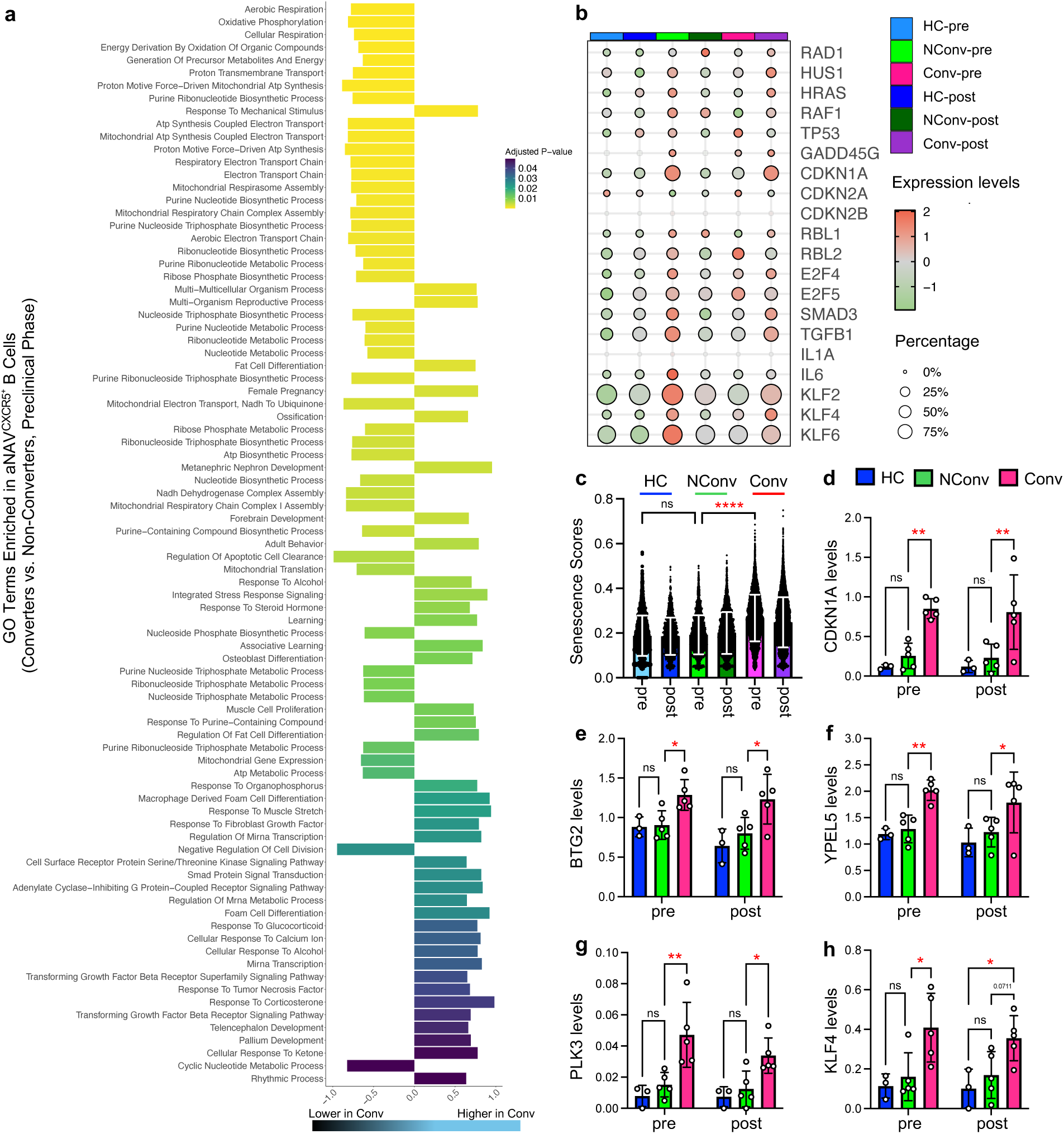
Converters’ aNAV^CXCR5⁺^ cells exhibit a senescent and developmentally arrested phenotype. (**a**) GO term enrichment analysis of aNAV^CXCR5⁺^ cells comparing Converters and Nonconverters at the preclinical stage. The x-axis indicates enrichment scores; values above 0.0 represent pathways enriched in Converters, and values below 0.0 represent pathways downregulated in Converters. Color scale denotes adjusted P values. (**b**) Dot plot showing expression of key senescence-associated genes, including upstream DNA damage and stress-response genes (HUS1, HRAS, RAF1), the senescence master regulator TP53 and CDKN1A, cell cycle arrest regulators (RBL1, RBL2, E2F4, E2F5), SASP factors (IL6, IL1A, TGFB1), and KLF family transcription factors. (**c**) Senescence scores calculated from a set of cell cycle arrest–related genes. (**d–h**) Patient-level expression of representative senescence regulators (CDKN1A, BTG2, YPEL5, PLK3, KLF4). Each dot represents an individual patient. *P < 0.05, **P < 0.01, ****P < 0.0001 by two-way ANOVA (c-h); ns, not significant. HC: ACPA^-^ Controls; NConv: ACPA^+^ Nonconverters; Conv: ACPA^+^ Converters; Pre: First time point (pre-clinical); Post: Second time point (post-onset).

A senescence score derived from a cell-cycle arrest gene set (Supplementary Table 2) was significantly higher in aNAV^CXCR5⁺^ cells from Converters than from Nonconverters and Controls at the pre-clinical RA time point and remained elevated after clinical RA (Fig. 3c). Key regulators, including CDKN1A, BTG2, YPEL5, PLK3, and the naive B-cell proliferation repressor KLF4, were consistently upregulated in all Converters at both time points (P < 0.05 versus Nonconverters and Controls) (Fig. 3d–h). Neither the senescence scores nor the expression of these genes differed significantly between Nonconverters and Controls, nor between the pre- and post-onset timepoints within each group. Thus, Converters’ aNAV^CXCR5⁺^ acquire a stable senescent-like, developmentally arrested state characterized by metabolic suppression, cell-cycle blockade, and a persistent SASP that emerges preclinically and endures through development of clinical RA.

### aNAV^CXCR5⁺^ skew the B-cell transcriptome toward pathogenic programs

Differential gene expression analysis identified the initial emergence of a pathogenic activation signature at the aNAV^CXCR5⁺^ stage in Converters at the preclinical RA time point. Top DEGs included activation (CD69, FOS, FOSB, JUN, JUND, NR4A2), feedback regulators (TSC22D3, NFKBIA, ZFP36), antigen presentation (HLA-DRB5), cellular stress (PPP1R15A), cytoskeletal/membrane remodeling (TUBA1A, MYADM), senescence-associated (CDKN1A, YPEL5, CSPNP1), and interferon-stimulated (IRF1) genes (Extended Data Fig. 1a).

A substantial portion of this activation–stress program persisted across differentiation in Converters. SwMem cells retained elevated CD69, NFKBIA, FOS/FOSB/JUND, KLF2, KLF6, DUSP1, PPP1R15A; ABCs retained high FOS, FOSB, JUND, YPEL5, KLF2, KLF6, DUSP1, PPP1R15A (Extended Data Fig. 1a). Several of these signatures remained detectable even after clinical onset (Extended Data Fig. 1b). Within Converters, comparisons between pre-clinical RA and clinical RA samples showed few DEGs across subclusters (Extended Data Fig. 1c), indicating that this program is established at the asymptomatic pre-clinical RA phase and remains largely stable through the development of clinical RA. These data suggest that aNAV^CXCR5⁺^ cells are in an early inflection point that initiates a durable, pathogenic transcriptional state.

### Converters’ aNAV^CXCR5⁺^ express unmutated IgM with distinct V–J signatures

VDJ-seq revealed preclinical alterations in the BCR repertoire of Converters. In Controls and Nonconverters, IgM accounted for ∼60–74% of BCRs; in Converters, it increased to 80% (pre-clinical RA) and 81% (clinical RA) (Fig. 4a). In Converters, IgM was predominantly restricted to aNAV^CXCR5⁺^ cells (Fig. 4b). Somatic hypermutation (SHM) was low in rNAV, aNAV^CXCR5⁺^, and aNAV^CD38hi^ across groups, rising in Transit and peaking in SwMem, without group differences (Fig. 4c). The activation score, calculated by a set of immune activation genes (Supplementary Table 3), was highest in Converters’ IgM-expressing aNAV^CXCR5⁺^ cells compared to their post-onset counterparts, Nonconverters, and Controls (P < 0.0001), indicating robust immune activation during the pre-clinical RA aNAV^CXCR5⁺^ stage in Converters (Fig. 4d). Converters also exhibited increased BCR repertoire diversity in aNAV^CXCR5+^ cells, with trends toward higher Shannon diversity and unique clone fractions at the pre-clinical RA phase, reaching significance at clinical RA phase (P < 0.05 versus both comparators) (Fig. 4e,f). Overall V and J gene usage was broadly similar, with modest shifts (IGHV6, IGHJ4, IGHJ6, IGLV2, IGLV3) (Extended Data Fig. 2). PCA of V–J rearrangements clearly separated Converters for both heavy and light chains (Fig. 4g), and heatmaps showed consistent enrichment of specific V–J combinations across all five Converters at both time points, remaining low in Nonconverters and Controls (Extended Data Fig. 3).

**Figure 4.**
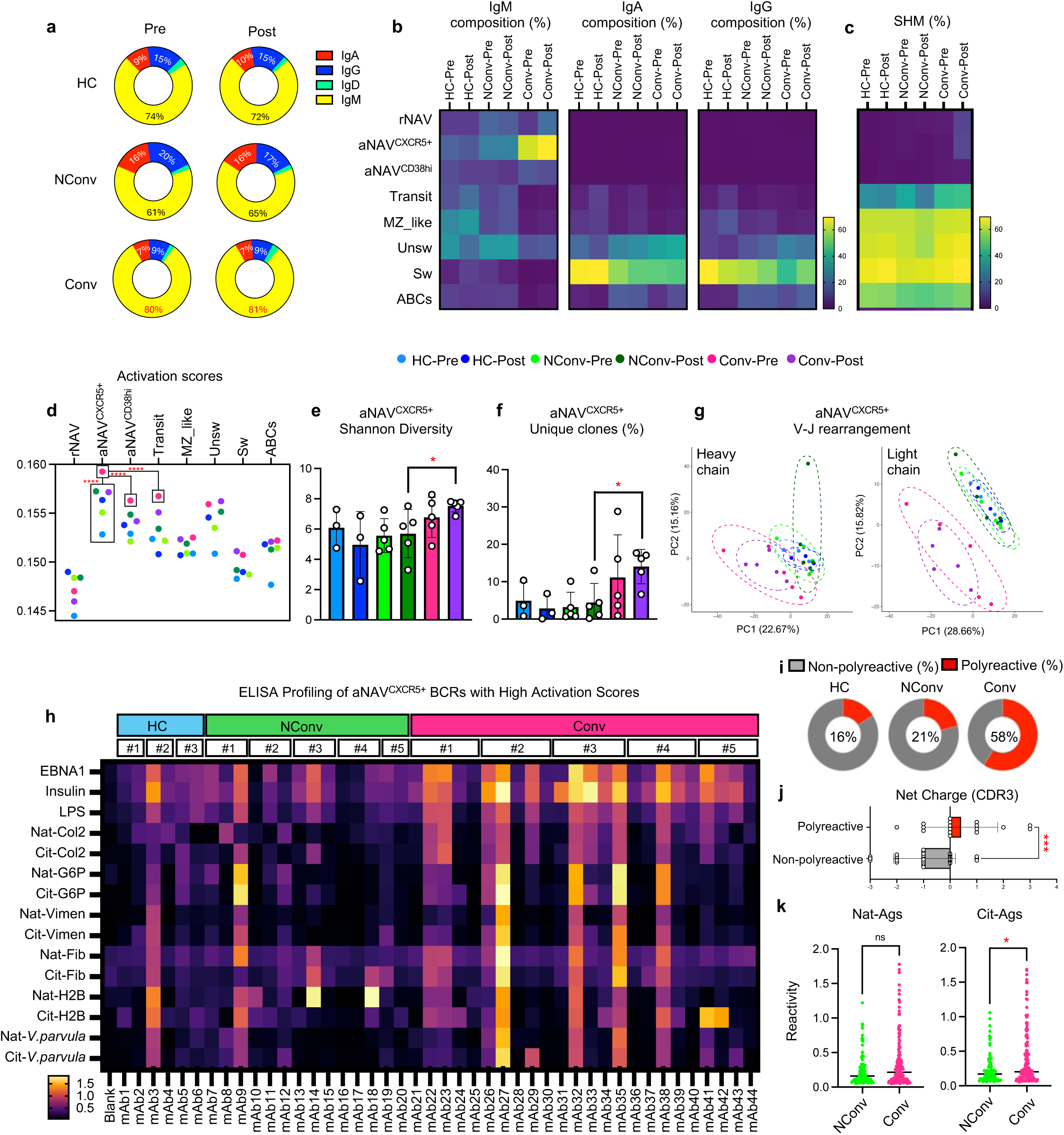
aNAV^CXCR5⁺^ cells express unmutated, polyreactive IgM and dominate the BCR repertoire in Converters. (**a**) Isotype composition of the BCR repertoire in Controls, Nonconverters, and Converters. (**b**) Distribution of IgM-, IgA-, and IgG-expressing BCRs across B cell subsets. For each isotype, all BCRs were pooled per condition, normalized to 100%, and the proportional contribution of each subset was calculated. (**c**) Proportion of somatically hypermutated BCRs across subsets. (**d**) Activation scores of BCR-expressing cells; cells without paired BCR data were excluded. Each dot represents the mean activation score of a given subset in one condition. (**e**) Shannon diversity index of aNAV^CXCR5⁺^ cells; each dot represents one patient. (**f**) Proportion of unique clones within the aNAV^CXCR5⁺^ repertoire; each dot represents one patient. (**g**) PCA of heavy-chain (left) and light-chain (right) V–J rearrangement patterns; each dot represents one patient. (**h**) Heatmap of ELISA reactivity for recombinantly expressed BCRs derived from IgM⁺ aNAV^CXCR5⁺^ cells with high activation scores at the pre-clinical RA phase, with one clone selected from each participant (#1–#5). Antigens tested included native (Nat) and citrullinated (Cit) collagen type II (Col2), glucose-6-phosphate isomerase (G6P), vimentin (Viment), fibrinogen (Fib), histone H2B (H2B), and bacterial lysates of *Veillonella parvula*. (**i**) Proportion of ELISA-confirmed polyreactive versus non-polyreactive mAbs. Experiments were performed in triplicate, and average optical density (OD) values were calculated. N = 6 for Controls, N = 14 for Nonconverters, and N = 24 for Converters. (**j**) Net CDR3 charge of polyreactive versus non-polyreactive BCRs. (**k**) ELISA reactivity of individual mAbs (10 µg/mL), showing a modest but significant increase in binding to citrullinated antigens in Converters compared to Nonconverters. No significant difference was observed in reactivity toward native antigens between the two groups. *P < 0.05, **P < 0.01; ***P < 0.001; ****P < 0.0001 by unpaired parametric Student’s t test (j), or by Kruskal–Wallis test (d), or by one-way ANOVA (e, and f), or by Mann-Whitney test (k); ns, not significant. HC: ACPA^-^ Controls; NConv: ACPA^+^ Nonconverters; Conv: ACPA^+^ Converters; Pre: First time point (pre-clinical); Post: Second time point (post-onset).

### Polyreactive IgM⁺ aNAV^CXCR5⁺^ recognize RA-associated autoantigens

To assess the antigen reactivity of aNAV^CXCR5⁺^ cells, we selected BCR sequences from this subset based on their activation scores, calculated using a defined set of immune activation genes (see Methods). These scores reflect transcriptional activation and are consistent with those shown in Fig. 4d. Specifically, we selected the top-ranking BCR sequences with the highest activation scores from Controls, Nonconverters, and Converters at the pre-clinical RA phase (Supplementary Table 4). These BCRs were recombinantly expressed as monoclonal antibodies (mAbs) using a human IgG backbone and tested by ELISA against a diverse antigen panel, including the Epstein–Barr virus antigen EBNA1 (a proposed autoimmune trigger [37, 38]); insulin; lipopolysaccharide (LPS); multiple RA-associated antigens in native and citrullinated forms (Collagen II, G6P, Vimentin, Fibrinogen, and H2B); and bacterial lysates of *Veillonella parvula*, an oral microbe implicated in RA pathogenesis [39, 40]. mAbs derived from Converters’ aNAV^CXCR5⁺^ cells exhibited broader and stronger reactivity across antigens compared with those from Nonconverters and Controls (Fig. 4h). Polyreactive BCRs were detected in all five Converters, with their frequency markedly increased (58%) relative to Nonconverters (21%) and Controls (16%) (Fig. 4i). ProtParam analysis revealed that ELISA-confirmed polyreactive mAbs possessed significantly higher net positive charge in their CDR3 regions compared to non-polyreactive mAbs (Fig. 4j). To further assess antigen binding strength, we selected representative polyreactive mAbs for ELISA-based titration against native and citrullinated RA-associated antigens. Polyreactive mAbs derived from Converters exhibited stronger binding across both antigen forms than those from Nonconverters (Extended Data Fig. 4). Notably, Converter-derived aNAV mAbs showed a modest but significant increase in binding affinity toward citrullinated antigens (Fig. 4k), while no overall difference was observed for native antigen binding. This preferential recognition of citrullinated antigens likely reflects the reduced positive charge caused by poly-citrullination, which enhances electrostatic complementarity with the positively charged CDR3 loops of these autoreactive antibodies.

### IgM⁺ aNAV^CXCR5⁺^ remain “stuck” through RA onset

While most aNAV^CXCR5⁺^ BCRs were singletons, clonal expansions were observed, with family members distributed within aNAV and across other B cell subsets (Fig. 5a). In Converters, singleton aNAV^CXCR5⁺^ cells had higher activation scores than those in Nonconverters or Controls (Fig. 5b). Whereas clonal aNAV^CXCR5⁺^ cells from Nonconverters and Controls showed lower activation scores than singletons, clonal aNAV^CXCR5⁺^ cells in Converters exhibited elevated activation scores comparable to singletons (Fig. 5b). Phylogenetic reconstruction showed frequent clustering of aNAV with non-aNAV relatives (“aNAV-related” cells) (Fig. 5c; Extended Data Fig. 5). aNAV-related clones were markedly increased in Converters (51.30 ± 6.68%) versus Nonconverters (11.33 ± 8.26%) and Controls (7.84 ± 5.45%) (P < 0.0001) (Fig. 5d).

**Figure 5.**
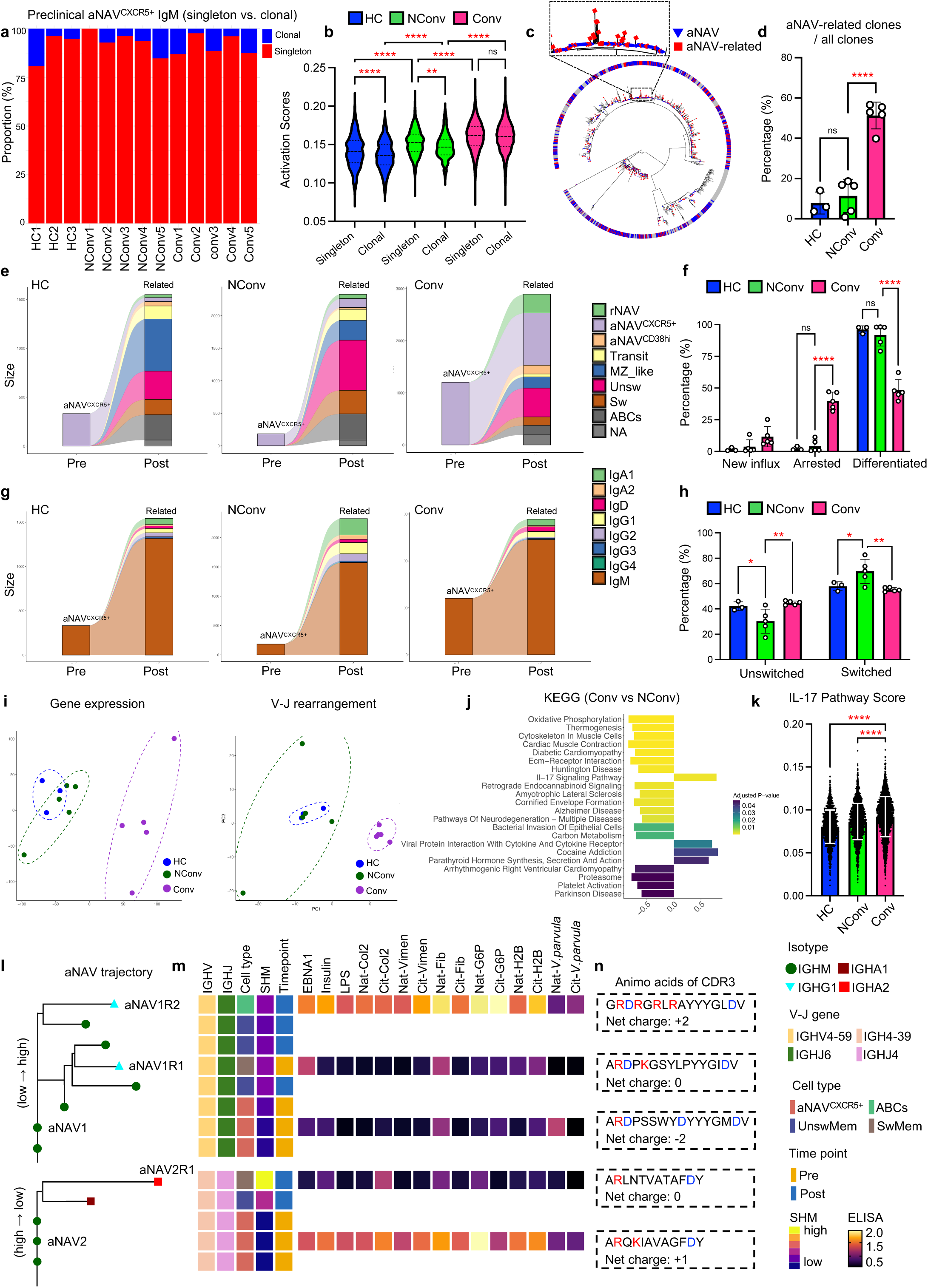
Distinct trajectory of aNAV^CXCR5⁺^ cells during RA onset. (**a**) Proportional analysis of singleton and clonal aNAV^CXCR5⁺^ IgM⁺ B cells at the pre-clinical RA phase for each participant. Singletons (red) represent unique BCRs with no clonal relatives; clonal aNAVs (blue) have at least one clonal family member based on heavy-chain V–J gene usage and CDR3 sequence similarity. (**b**) Violin plots showing activation scores of singleton and clonal aNAV^CXCR5⁺^ cells across Controls, Nonconverters, and Converters at pre-clinical RA phase. (**c**) Representative phylogenetic tree illustrating the evolutionary relationship between aNAV and aNAV-related BCRs. Full trees for all clones are shown in Supplementary Figure 5. (**d**) Percentage of aNAV-related clones (defined as clones containing at least one aNAV^CXCR5⁺^ BCR) among all clones (across pre- and post-onset time points) for each patient. (**e**) IgM⁺ aNAV^CXCR5⁺^ cells at the pre-clinical phase were subsetted, and their clonal relatives were identified at the post-onset stage; schematic shows cell type transitions from preclinical aNAV^CXCR5⁺^ cells to post-onset aNAV-related cells. (**f**) Percentages of new influx (rNAV), arrested (aNAV^CXCR5⁺^), and differentiated (later-phase B cell subsets) aNAV-related cells at post-onset. Each dot represents one individual. (**g**) Isotype trajectories from preclinical IgM⁺ aNAV^CXCR5⁺^ cells to post-onset aNAV-related BCRs. (**h**) Percentages of unswitched and class-switched BCRs among post-onset aNAV-related BCRs for each patient. (**i**) PCA of transcriptional profiles (left) and V–J rearrangement patterns (right) showing distinct post-onset aNAV-related cells in Converters compared with Nonconverters and Controls. (**j**) KEGG pathway analysis indicating significant enrichment of IL-17 signaling in post-onset aNAV-related cells from Converters. (**k**) IL-17 signaling pathway scores across groups. (**l**) Reactivity trajectory analysis of aNAV clonal families: IgM⁺ BCRs from Converter aNAV^CXCR5⁺^ cells at the preclinical stage and class-switched, somatically hypermutated aNAV-related BCRs at the post-onset stage were recombinantly expressed and tested by ELISA. For each antibody, IGHV/IGHJ gene usage, cell type, SHM rate, and timepoint are indicated. Reactivity to multiple antigens—including EBNA1, insulin, LPS, and native (Nat) or citrullinated (Cit) RA antigens—is shown. Experiments were performed in triplicate, and average OD values were calculated. (**m**) Corresponding CDR3 amino acid sequences with calculated net charges; positively charged residues are in red, negatively charged residues in blue. *P < 0.05; **P < 0.01; ****P < 0.0001 by one-way ANOVA (d and k), or by two-way ANOVA (b, f, and h); ns, not significant. HC: ACPA^-^ Controls; NConv: ACPA^+^ Nonconverters; Conv: ACPA^+^ Converters.

Clonal tracing of pre-clinical IgM⁺ aNAV^CXCR5⁺^ showed that in Nonconverters and Controls, most progressed into aNAV^CD38hi^, Transit, MZ-like, UnswMem, SwMem, and ABCs (Fig. 5e,f). In Converters, a substantial fraction remained in early aNAV across both time points, consistent with senescence-associated developmental arrest. This arrested fraction expanded across all Converters, with fewer differentiated progeny.

Post-onset of clinical RA, the frequency of new aNAV-related rNAV cells trended higher in Converters than in Nonconverters and Controls (Fig. 5e,f), suggesting ongoing influx of polyreactive naive B cells from bone marrow to periphery. No differences in influx/arrested/differentiated fractions were observed between Nonconverters and Controls. Isotype tracking showed that, across all groups, aNAV^CXCR5⁺^ IgM largely remained IgM from the first to the second time point, with occasional switching to IgA/IgG (Fig. 5g). Notably, class-switched aNAV-related cells were more frequent in Nonconverters than in Converters or Controls (Fig. 5h), suggesting a distinct class-switch trajectory in ACPA⁺ non-converters. PCA indicated that post-onset aNAV-related cells from Converters had distinct transcriptomic and V–J profiles (Fig. 5i). KEGG analysis showed upregulated IL-17 signaling and viral protein interaction pathways and downregulated oxidative phosphorylation (Fig. 5j). Consistently, IL-17 pathway scores (Supplementary Table 5) were significantly higher in Converters’ aNAV-related cells (Fig. 5k), indicating that the metabolic/inflammatory state established in pre-clinical RA aNAV^CXCR5⁺^ is propagated to clonal relatives at onset.

### Divergent reactivity trajectories of aNAV^CXCR5⁺^-derived BCRs during RA onset

To assess functional evolution, we expressed antibodies from IgM⁺ aNAV cells at the pre-clinical RA phase and from their class-switched, somatically mutated relatives at the clinical RA phase from the same Converter patient. Between the pre-clinical and clinical phases, two distinct trajectories of antigen reactivity emerged (Fig. 5l–n). In the “low-to-high” reactivity trajectory, SHM progressively broadened antigen reactivity and increased binding affinity. For the aNAV1 family, the mutated IgG1 antibody aNAV1R2 (derived from ABCs at the clinical RA stage) showed markedly broader polyreactivity, spanning insulin, LPS, EBNA1, and multiple native and citrullinated RA antigens, compared with its precursors. Across this family, the CDR3 net charge progressed from −2 in aNAV1, to 0 in aNAV1R1, and to +2 in aNAV1R2. In the “high-to-low” reactivity trajectory, SHM attenuated polyreactivity. In the aNAV2 family, the mutated IgA2 antibody aNAV2R1 (from switched memory cells at the clinical RA stage) displayed reduced polyreactivity relative to the original aNAV2, accompanied by a shift in CDR3 net charge from +1 to 0. Thus, although many polyreactive IgM⁺ aNAVs persist through disease onset, somatic hypermutation and class switching can either amplify or attenuate their polyreactivity in antibody-secreting descendants, thereby reshaping the autoreactive landscape during RA conversion.

### Polyreactive IgM⁺ aNAV^CXCR5⁺^ infiltrate the RA synovium

We next assessed whether aNAV^CXCR5⁺^ localize to inflamed joints. Synovial tissues were collected from an independent cohort that consisted of three individuals with ACPA⁺ RA and four with ACPA⁻ OA; Supplementary Table 6). Synovial B cells were enriched and profiled by integrated 5’ scRNA-seq/VDJ/CITE-seq and combined with a published RA/OA synovial dataset, yielding nine RA and seven OA samples (Fig. 6a,b). Among 26,290 CD19⁺ synovial B cells (Fig. 6c), UMAP identified a plasma/plasmablast cluster and multiple B-cell populations, including a discrete aNAV cluster (Fig. 6d). Synovial aNAVs mirrored the peripheral phenotype (CD19⁺ CD27⁻ IGHM⁺ IGHD⁺ CXCR5⁺) with high CD69 (Fig. 6e). Among B-cell subsets, synovial aNAV^CXCR5⁺^ expressed the highest HLA-DRA, suggesting strong antigen-presenting capacity. Quantitatively, aNAV^CXCR5⁺^ cells were expanded in RA vs OA (Fig. 6f). GO analysis showed enrichment of B-cell–mediated immune pathways in RA aNAV^CXCR5⁺^ (Fig. 6g). Most synovial aNAV BCRs remained IgM⁺ with low levels of somatic hypermutation in both RA and OA (Fig. 6h,i), but RA aNAV^CXCR5⁺^ tended toward higher repertoire diversity (RA: 2.98 ± 1.26 vs. OA: 1.41 ± 1.18) and unique clones (RA: 30.4 ± 37.3% vs. OA: 6.62 ± 9.26%) (Fig. 6j,k). ELISA profiling detected polyreactive BCRs (Supplementary Table 7) among synovial aNAV^CXCR5⁺^ in both RA and OA, with higher frequency in RA (42%, n=12) versus OA (27%, n=11) (Fig. 6l,m).

**Figure 6.**
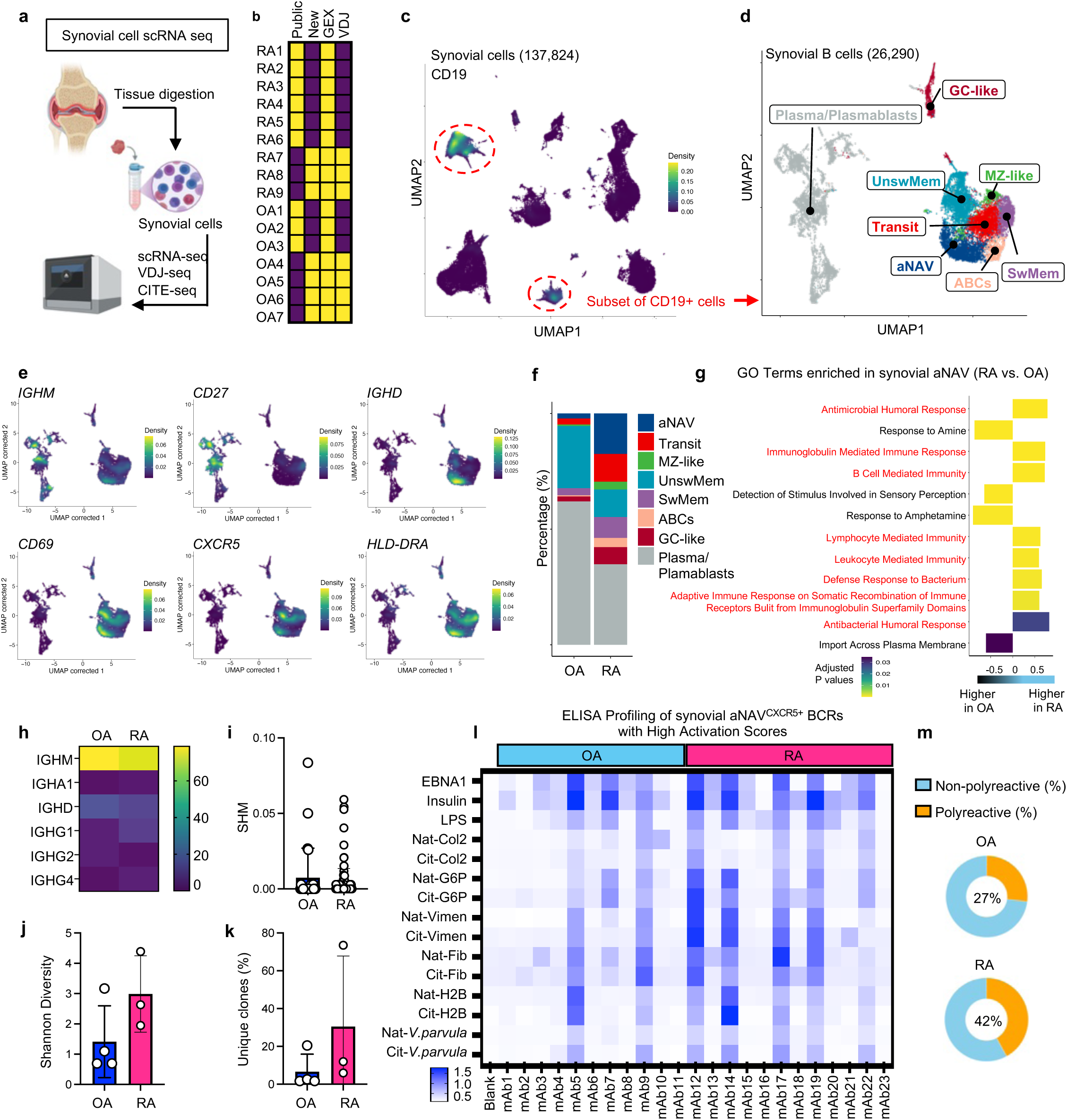
Polyreactive aNAV^CXCR5⁺^ cells are present and expanded in RA synovium. (**a**) Experimental workflow for synovial tissue processing. Synovial samples from RA (n = 3) and OA (n = 4) patients were enzymatically digested, followed by B cell enrichment, scRNA-seq, VDJ-seq, and CITE-seq. (**b**) Newly generated datasets were integrated with a published synovial scRNA-seq dataset (6 RA and 4 OA patients), yielding a combined cohort of 9 RA and 7 OA samples. (**c**) CD19⁺ B cell clusters were identified from synovial single-cell data and subsetted for downstream analysis. (**d**) UMAP visualization of synovial B cells (26,290 cells), showing major subsets: plasma/plasmablasts, aNAV, transitional, unswitched memory (UnswMem), marginal zone-like (MZ-like), switched memory (SwMem), and germinal center-like (GC-like) compartments. (**e**) Density plots of key aNAV^CXCR5⁺^ cell markers, including IGHM, CD27, IGHD, TCL1A, CD69, CXCR5, and HLA-DRA. (**f**) Proportional composition of B cell subsets in RA versus OA synovium, showing significant expansion of aNAV^CXCR5⁺^ cells in RA. (**g**) GO term enrichment comparing aNAV^CXCR5⁺^ cells between RA and OA synovium. Positive enrichment scores indicate RA-enriched pathways; negative scores indicate OA-enriched pathways. (**h**) Isotype distribution of aNAV^CXCR5⁺^ cells in RA and OA synovium. (**i**) Somatic hypermutation (SHM) rates of aNAV^CXCR5⁺^ BCRs in RA and OA. (**j**) Shannon diversity index of aNAV^CXCR5⁺^ repertoires in RA and OA. (**k**) Proportion of unique clones among aNAV^CXCR5⁺^ cells in RA and OA. For transcriptomic analyses, n = 9 RA and n = 7 OA; for VDJ repertoire analyses, n = 3 RA and n = 4 OA. (**l**) ELISA assays of recombinant mAbs derived from aNAV^CXCR5⁺^ cells with high activation scores, selected from both OA and RA patients. Experiments were performed in triplicate, and average OD values were calculated. (**m**) Percentage of polyreactive versus non-polyreactive BCRs in each group. N = 11 for OA BCRs and N = 12 for RA BCRs.

### Citrullinated antigens and CpG act as dual triggers of naive-B-cell activation and senescence in RA

Given the limited availability of pre-clinical RA samples, we next investigated whether aNAV populations could also be detected in an independent cohort of patients with clinically established ACPA⁺ RA (Supplementary Table 8). To this end, we developed a flow cytometry strategy to identify and isolate aNAV cells from PBMCs of ACPA⁺ RA patients, using the markers CD19, CD27, IgM, IgD, CXCR5, and CD69 (Fig. 7a). Flow cytometry showed increased frequencies of CD69⁺ single-positive and CD69⁺CXCR5⁺ double-positive cells within the CD27⁻ IgM⁺ IgD⁺ naive compartment in ACPA⁺ RA compared with healthy controls (Fig. 7b,c), with no significant changes in CD27⁺ memory or double-negative B cells (Fig. 7d–g).

**Figure 7.**
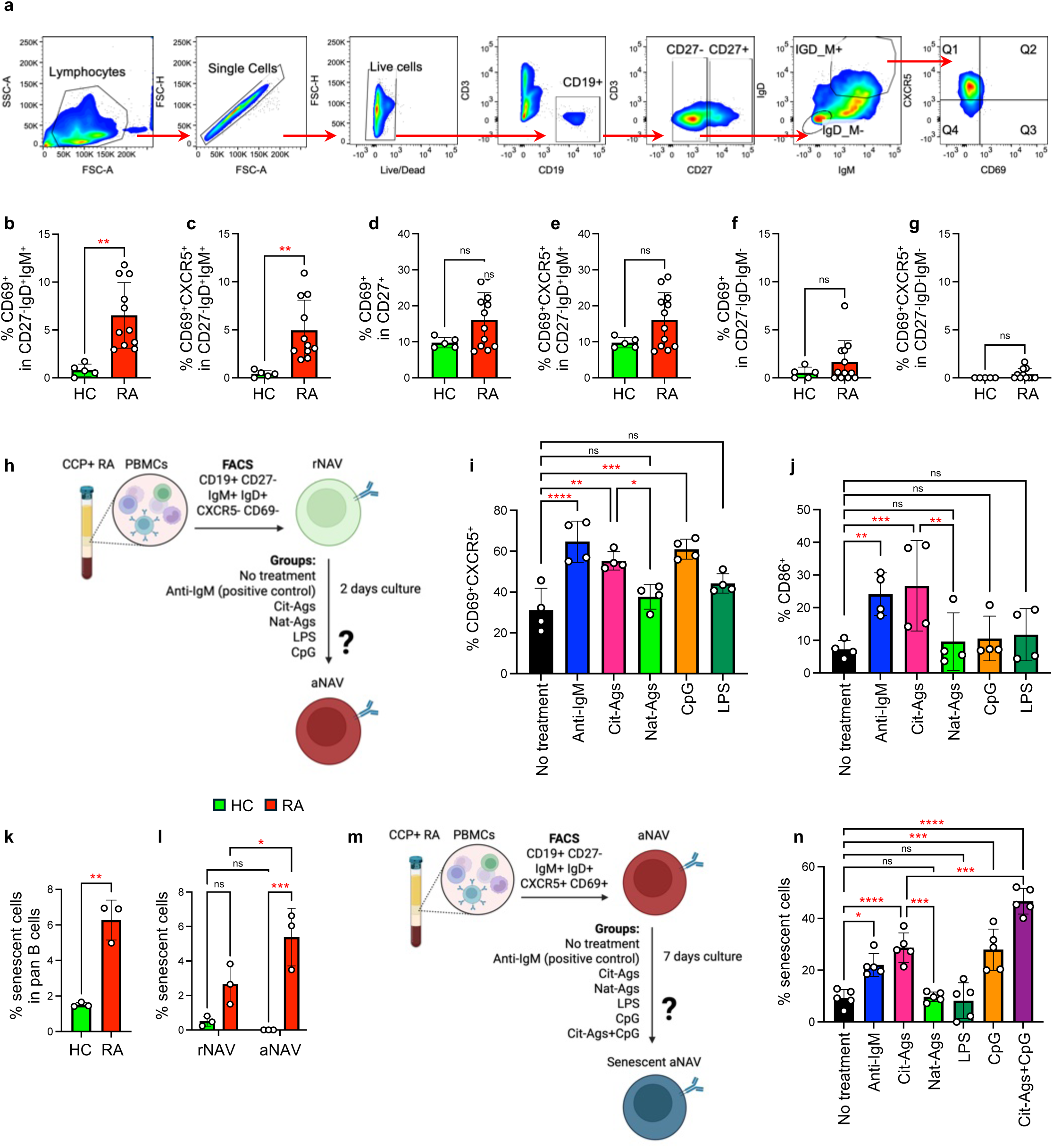
Effects of antigen exposure on naive B cell activation and senescence in RA. (**a**) Gating strategy for FACS sorting of CD19⁺CD27⁻IgM⁺IgD⁺CXCR5⁺CD69⁺ aNAV cells. (**b–g**) Frequencies of CD69⁺ or CD69⁺CXCR5⁺ cells across naive, memory, and double-negative B cell subsets in peripheral blood from ACPA^+^ RA patients and healthy controls. (**h**) Experimental design for antigen screening to identify potential triggers of naive B cell activation. rNAV cells were FACS-sorted and cultured for 2 days with anti-IgM, citrullinated antigens (Cit-Ags), native antigens (Nat-Ags), CpG, or LPS. (**i**) Percentages of CD69⁺CXCR5⁺ cells after stimulation. N = 4 per group. (**j**) Percentages of CD86⁺ cells after stimulation. N = 4 per group. (**k**) Frequencies of senescent cells in total B cells from ACPA^+^ RA patients and healthy controls. N = 3 per group. (**l**) Frequencies of senescent cells in rNAV and aNAV subsets from ACPA^+^ RA patients and healthy controls. N = 3 per group. (**m**) Experimental design for antigen screening to identify potential inducers of aNAV senescence. aNAV cells were FACS-sorted and cultured for 7 days under stimulation with anti-IgM, Cit-Ags, Nat-Ags, CpG, LPS, or Cit-Ags + CpG. (**n**) Percentages of senescent cells after stimulation. N = 5 per group. *P < 0.05; **P < 0.01; ***P < 0.001; ****P < 0.0001 by Student’s t test (b-g and k), or by one-way ANOVA (i, j, and n), or two-way ANOVA (l); ns, not significant.

To test whether antigens drive rNAV-to-aNAV activation, sorted rNAV (CD19⁺ CD27⁻ IgM⁺ IgD⁺ CXCR5⁻ CD69⁻) were stimulated for 2 days with native/citrullinated RA antigens (histone H3, vimentin, fibrinogen), LPS, or CpG (TLR9 agonist) (Fig. 7h). Anti-IgM, a broad BCR stimulator, served as a positive control. Citrullinated antigens or CpG robustly induced rNAV activation (increased CD69⁺CXCR5⁺) comparable to anti-IgM; native antigens and LPS did not (Fig. 7i). Moreover, citrullinated antigens, but not CpG/native/LPS, significantly increased CD86⁺ rNAVs (Fig. 7j), indicating an antigen-specific enhancement of co-stimulatory capacity.

We next assessed senescence by measuring activation of β-galactosidase, a well-known marker for cellular senescence [41]. Overall, ACPA⁺ RA showed higher B-cell senescence than healthy controls (Fig. 7k). Both rNAV and aNAV had elevated senescence in RA, with aNAV highest (HC: rNAV 0.50 ± 0.28%, aNAV 0 ± 0%; RA: rNAV 2.65 ± 1.16%, aNAV 5.38 ± 1.66%; P < 0.05 for RA aNAV vs all) (Fig. 7l). To model chronic antigen stimulation, sorted aNAV (CD19⁺ CD27⁻ IgM⁺ IgD⁺ CXCR5⁺ CD69⁺) from ACPA⁺ RA were stimulated for 7 days with native/citrullinated antigens, LPS, CpG, or CpG + citrullinated antigens (Fig. 7m). Citrullinated antigens and CpG each induced aNAV senescence, whereas native antigens and LPS had minimal effects; CpG + citrullinated antigens produced a synergistic increase beyond either alone (P < 0.001) (Fig. 7n). Collectively, these findings indicate that dual stimulation via BCR engagement (by citrullinated autoantigens) and TLR9 signaling (by CpG) drives both activation and senescence of naive B cells in RA.

### aNAVs activate CD4⁺ and CD8⁺ T cells via HLA-dependent presentation of citrullinated antigens

Because aNAVs are not antibody-secreting cells, we hypothesized they contribute to RA via antigen presentation. We sorted rNAV, aNAV, CD27⁺ memory B cells, and CD3⁺ T cells from ACPA⁺ RA PBMCs (Fig. 8a). B cells were pre-stimulated with native or citrullinated RA antigens (anti-IgM as positive control) and co-cultured with autologous T cells (1:10) for 7 days, with or without HLA class I/II blockade.

**Figure 8.**
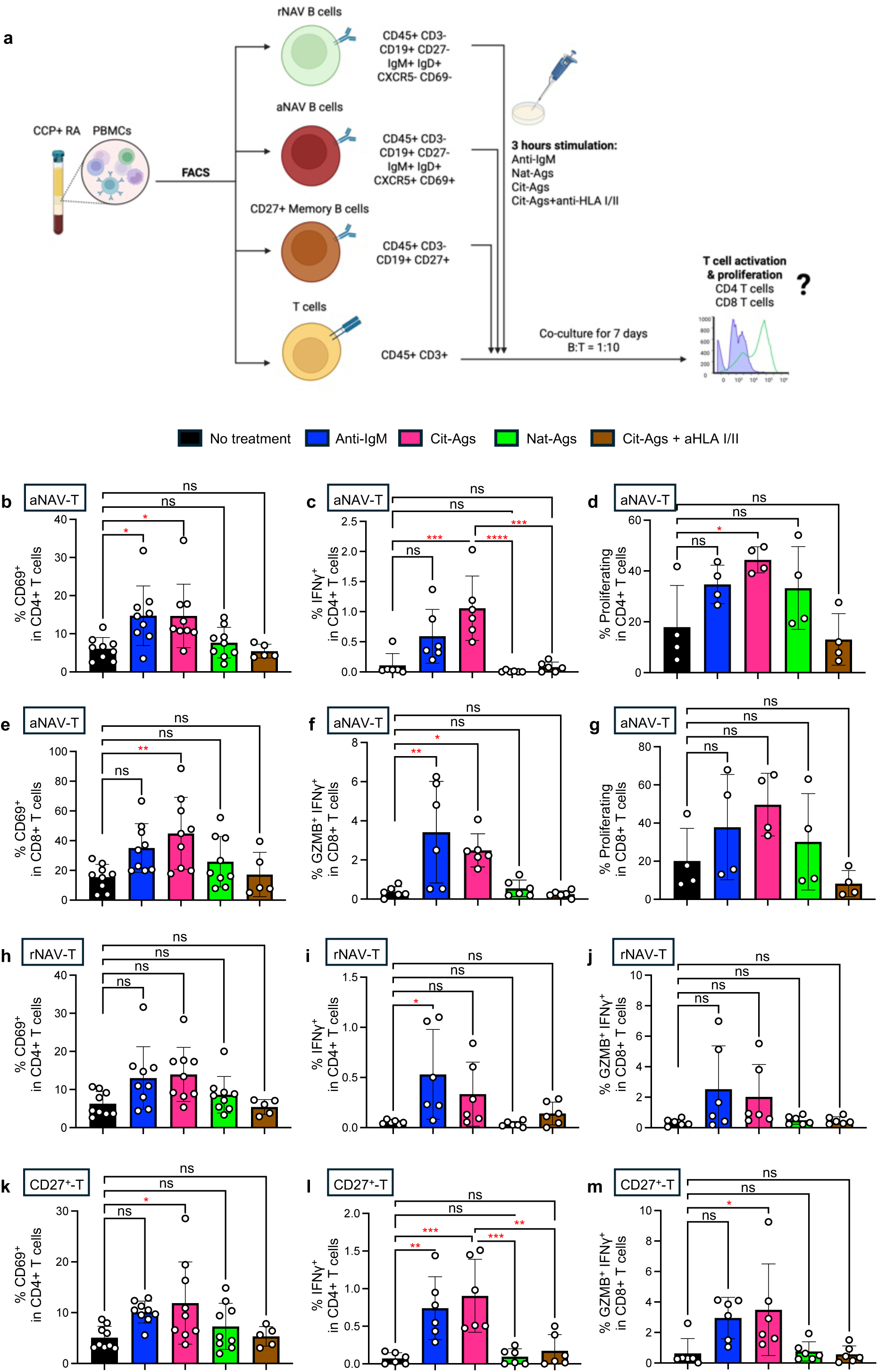
Citrullinated antigen–stimulated aNAV cells activate T cells via antigen presentation. (**a**) Experimental design to assess whether antigen-stimulated B cells activate T cells. rNAV, aNAV, and CD27⁺ memory B cells were FACS-sorted from ACPA^+^ RA patients based on indicated markers, stimulated with antigens for 3 h, and then co-cultured with CD3⁺ T cells for 7 days. T cell activation and proliferation were analyzed by flow cytometry. (**b-d**) Percentages of CD69⁺, IFNγ⁺, and proliferating CD4⁺ T cells co-cultured with aNAV cells. N = 4–9 per group. (**e-g**) Percentages of CD69⁺, GZMB⁺IFNγ⁺, and proliferating CD8⁺ T cells co-cultured with aNAV cells. N = 4–9 per group. (**h-i**) Percentages of CD69⁺ and IFNγ⁺ cells among CD4⁺ T cells co-cultured with rNAV cells. N = 5–9 per group. (**j**) Percentage of GZMB⁺IFNγ⁺ cells among CD8⁺ T cells co-cultured with rNAV cells. N = 6 per group. (**k-l**) Percentages of CD69⁺ and IFNγ⁺ CD4⁺ T cells co-cultured with CD27⁺ memory B cells. N = 5–9 per group. (**m**) Percentage of CD69⁺IFNγ⁺ CD8⁺ T cells co-cultured with CD27⁺ memory B cells. N = 6 per group. *P < 0.05; **P < 0.01; ***P < 0.001; ****P < 0.0001 by one-way ANOVA (b-m); ns, not significant.

aNAVs when loaded with citrullinated antigens robustly activate both CD4⁺ and CD8⁺ T cells (increased CD4⁺CD69⁺, CD4⁺IFN-γ⁺, CD8⁺CD69⁺, CD8⁺GZMB⁺IFN-γ⁺) and markedly enhanced proliferation, exceeding anti-IgM (Fig. 8a–g). In contrast, rNAVs did not activate CD4⁺ or CD8⁺ T cells after native or citrullinated antigen exposure (Fig. 8h– j). HLA class I/II blockade abrogated the ability of aNAVs + citrullinated antigens to promote CD4+ and CD8+ T cell activation, confirming presentation-dependent activation. Native antigen–stimulated aNAVs had minimal activity. CD27⁺ memory B cells also activated T cells upon citrullinated antigen exposure (Fig. 8k–m). Thus, presentation-driven T-cell activation emerges at the aNAV stage and is retained in memory progeny.

## Discussion

RA is increasingly recognized as a disease that develops and can be identified during an at-risk stage, with autoantibodies such as ACPAs appearing years before the onset of clinically-apparent inflammatory arthritis (i.e. clinical RA) [42, 43]. As a result, the field is shifting from reactive treatment after onset toward early diagnosis and preventive intervention. Although large cohort studies have revealed molecular differences between ACPA⁺ at-risk individuals and those with early RA [30, 32, 44–46], the specific cellular and molecular mechanisms driving progression remain incompletely defined. Here, by directly comparing ACPA⁺ Converters with age- and sex-matched Nonconverters, we identify a striking preclinical expansion of CXCR5⁺ CD69⁺ activated naive (aNAV) B cells in Converters that persists through clinical onset. These aNAV B cells are pro-inflammatory, senescence-prone, and developmentally arrested. They display distinct V–J recombination patterns and generate polyreactive, autoreactive BCRs capable of binding multiple RA-associated autoantigens. Moreover, aNAV expansion reshapes the overall BCR repertoire and skews the global B-cell transcriptome toward pathogenic programs. Functionally, aNAV B cells can infiltrate RA synovium and serve as APCs that present citrullinated antigens to activate both CD4⁺ and CD8⁺ T cells. Together, these findings establish a mechanistic framework in which preclinical aNAV expansion initiates autoimmune activation and drives RA onset. This newly defined pathway, coupled with the emerging success of anti-CD19 CAR-T therapies in autoimmune diseases, highlights testable strategies for B cell–targeted prevention prior to clinical disease manifestation.

The expanded aNAV cluster is phenotypically defined by CD19⁺ CD27⁻ IgM⁺ IgD⁺ CXCR5⁺ CD69⁺ expression. However, CXCR5 is not unique to aNAVs; it is also expressed by transitional B cells, marginal zone (MZ)-like B cells, and unswitched memory B cells [47]. CXCR5 is a chemokine receptor that binds CXCL13 and guides B cell migration toward germinal centers (GCs). In aNAVs, CXCR5 expression suggests prior antigen encounter and an attempt to localize to GCs for further selection and maturation [48, 49]. Within GCs, CXCR5⁺ B cells receive help from follicular helper T (Tfh) cells, undergo somatic hypermutation and class-switch recombination, and ultimately differentiate into high-affinity memory B cells or antibody-secreting plasma cells [50, 51]. Upon completing this program, CXCR5 is downregulated, permitting B-cell exit from the GC to peripheral sites [52]. GC-independent maturation can also occur in inflamed tissues, where peripheral helper T (Tph) cells provide alternative B cell help [53, 54]. In ACPA^-^ Controls and ACPA⁺ Nonconverters, CXCR5⁺ aNAVs appear to follow these canonical or alternative maturation pathways and successfully transition into later developmental stages. In contrast, ACPA⁺ Converters accumulate activated yet senescent and developmentally arrested CXCR5⁺ aNAVs, suggesting that both Tfh- and Tph-mediated maturation signals fail at the preclinical stage. While a subset of these aNAVs may still transition into memory-like or ABCs, the majority remain functionally stalled. This results in a “stuck” phenotype characterized by prolonged circulation, impaired differentiation, and persistent presence in peripheral blood or tissues throughout RA conversion.

Our study suggests that this “stuck” phenotype is driven by dysregulated B-cell signaling shaped by chronic antigenic stimulation, anergy-like functional tuning, and metabolic reprogramming, rather than tolerance breakdown alone. Notably, we observed upregulation of several Krüppel-like factor (KLF) family members—KLF2, KLF4, and KLF6—in preclinical aNAVs from Converters. These transcription factors, best known as tumor suppressors, may promote cell cycle arrest and cellular senescence, providing a mechanistic link to the senescence-like state of these arrested B cells [55–58]. Beyond enforcing arrest, KLFs are increasingly recognized for roles in B-cell fate. For example, KLF2 maintains a circulation-competent pre-GC phenotype by repressing GC entry [59, 60]; its elevated expression in Converters suggests that aNAVs are transcriptionally locked in a pre-GC state, preventing further differentiation and enabling persistence of autoreactive clones. KLF4, normally downregulated upon B-cell activation, acts as a brake on cell cycle progression; its sustained expression may reinforce quiescence or senescence [61–63]. KLF6, though less studied, is upregulated in activation-prone B cells in celiac disease and co-expressed with CD69, FOS, and JUN, a transcriptional signature also enriched in Converter aNAVs [64]. Together, these data point to KLF-mediated transcriptional reprogramming as a potential mechanism of aNAV arrest.

Immune aging refers to the progressive decline of immune function with age and is increasingly implicated in the pathogenesis of autoimmune diseases [65, 66]. This process involves widespread alterations in immune homeostasis, including heightened antigenicity of self-antigens and dysregulation of lymphocyte populations and immune repertoires. Together, these changes increase susceptibility to immune dysregulation and autoimmunity. In B cells, aging-associated senescence drives the emergence of a distinct subset known as age-associated B cells [67]. Although the precise mechanisms linking B cell senescence to autoimmunity remain incompletely understood, recent studies offer important insights. Continuous BCR signaling has been shown to promote the differentiation of age-associated B cells from anergic B cells in both aged and autoimmune-prone mice [68]. In parallel, persistent stimulation by commensal microbiota can induce senescence in germinal center B cells, thereby contributing to age-related alterations in oral and gut microbiota [69, 70]. These findings suggest that chronic antigenic stimulation stresses B cells, promotes senescence, and disrupts immune homeostasis. Furthermore, Viglianti and colleagues demonstrated that the activation of autoreactive B cells by CpG DNA requires sequential engagement of the BCR and TLR9 [71]. In this process, the BCR delivers DNA-containing immune complexes to endosomes, where TLR9 detects CpG motifs within the DNA. Consistent with this model, our study demonstrates that citrullinated self-antigens and CpG can synergistically induce senescence in naive B cells during their early activated phase. Because CpG motifs mimic viral DNA, these results implicate viral exposure as a potential trigger of B cell senescence and subsequent autoimmune activation. Notably, citrullinated antigens elicited a stronger senescence response than their native counterparts, although the molecular mechanisms by which citrullination enhances this effect remain to be elucidated. Together, these results support a model in which chronic antigen exposure promotes B cell senescence and contributes to autoimmune progression. They further suggest that citrullination of self-antigens may represent a critical factor in accelerating immune aging and predisposing to autoimmunity.

The presence of polyreactive and autoreactive naive B cells has been described in multiple autoimmune diseases, including RA [13, 33, 72–75]. This is thought to reflect failure of central and peripheral tolerance mechanisms that normally delete or suppress autoreactive clones. In addition, some autoreactive B cells enter anergy, a state that silences their differentiation into antibody-secreting cells [35]. In Converters, however, both deletion and regulation appear impaired. Pathogenic aNAVs exhibit enhanced downstream pro-inflammatory signaling despite attenuated proximal BCR signaling, a hallmark of anergy-like tuning. This is further supported by their developmental arrest phenotype, a defining feature of anergy-like B cells. Because aNAVs remain naive and unswitched, they do not secrete soluble antibodies, but they continue to express membrane-bound polyreactive IgM, enabling antigen recognition, presentation and T–B cell crosstalk. Continuous influx of naive B cells from the bone marrow likely contributes to the pool of aNAVs, but it is the peripheral arrest upon antigen encounter that drives their accumulation.

Although T cells have long thought to be central to the pathogenesis of RA and the strongest genetic risk factors for RA map to genetic polymorphisms in MHC class I and class I MHC genes which present antigens to T cells [76], a major unanswered question in the field has been how these autoreactive T cells become activated. Here, we demonstrate that aNAV B cells serve as APCs that uptake and present citrullinated antigens to drive anti-citrullinated antigen CD4 and CD8 T cell responses. In our previous work, we showed that citrullinated antigens promote the expansion of cytotoxic GZMB⁺ CD8⁺ T cells [77]. Consistently, Aslam et al. reported the emergence of pro-inflammatory autoreactive T cell responses in ACPA⁺ at-risk individuals [78]. Here, we extend these findings by showing that such T cell activation may be amplified at the pre-clinical RA phase through aNAV-mediated presentation of citrullinated antigens, establishing a cycle of reciprocal B–T cell activation that precedes symptom onset.

Polyreactive BCRs play a central role in this process. Under normal conditions, low-affinity polyreactivity is tolerated and broadens pathogen-specific binding, whereas high-affinity polyreactive BCRs are removed by deletion or receptor editing [79]. Although V and J gene usage was broadly similar between Converters and Nonconverters, their V–J recombination patterns were distinct, consistent with antigen-driven selective pressure. These skewed recombination patterns increase BCR diversity, enabling Converters’ aNAV-derived IgM to recognize a broader range of self-antigens, yielding more frequent polyreactive BCRs. Polyreactivity is thought to be governed primarily by the heavy-chain CDR3 [80]. Rabia et al. showed that positively charged CDRs increase risk of low specificity [81], consistent with our finding that polyreactive aNAV BCRs carry a significantly higher net positive charge in their heavy-chain CDR3 compared with non-polyreactive mAbs. Additional features, including CDR3 length, conformational flexibility, and germline usage and rearrangement, also contribute [82]. Moreover, aNAV-derived mAbs from Converters showed modestly higher binding to citrullinated RA antigens than to native forms. This preference is likely facilitated by the loss of positive charges on antigens during polycitrullination, which enhances electrostatic complementarity with positively charged CDR3s [83]. In line with this, we previously demonstrated that plasmablast-derived mAbs from ACPA⁺ RA patients frequently bound multiple citrullinated antigens, reflecting their polyreactive nature [84]. Importantly, from the preclinical to clinical RA phase, clonally related aNAV BCRs exhibited divergent trajectories during somatic hypermutation and class switching, leading to either increased or reduced polyreactivity in their post-onset antibody-secreting progeny. Thus, somatic diversification from aNAV IgM clones may drive epitope spreading and shape the functional evolution of autoreactive B cell lineages during RA progression.

While the anti-CCP assay is the most widely used diagnostic test for RA, autoantibodies against native antigens have also been reported. König et al. found that anti-native RA33 antibodies were enriched in early RA and associated with low erosion scores, whereas anti-citrullinated RA33 correlated with disease duration and erosive disease [85]. These findings suggest that autoreactivity to native antigens may precede that to citrullinated antigens, with citrullination amplifying pathogenicity at mucosal sites [25, 40]. Poulsen et al. similarly reported anti-native autoantibodies in both ACPA⁺ and ACPA⁻ RA patients [86]. In line with these observations, we found that aNAV-derived IgM could bind both native and citrullinated RA antigens, but only citrullinated antigens effectively induced activation, senescence, and T-cell stimulation. Native antigens were bound but had minimal stimulatory impact. Moreover, citrullinated antigens combined with CpG, a TLR agonist, synergistically amplified the senescence phenotype of aNAVs, suggesting cooperation between BCR and innate signaling in pathogenic reprogramming. Moreover, IgM-expressing aNAV^CXCR5⁺^ cells exhibited higher activation scores than other B cells at the pre-clinical RA phase, and even higher than aNAV^CXCR5⁺^ cells at clinical RA phase, indicating that these early aNAVs undergo pronounced antigen-driven immune activation during the initial phase of disease evolution, well before clinical onset. Because of their anergy-like state, pathogenic aNAVs predominantly express membrane-bound polyreactive IgM, which is not detected by current serological assays. Developing new diagnostic approaches to identify such membrane-bound autoreactive IgMs in the blood of ACPA⁺ at-risk individuals will be critical for improving early detection and prevention strategies.

Researchers have made significant progress in defining the molecular landscape of the ACPA⁺ at-risk phase, underscoring the immune activation precedes the onset of clinical RA. Single-cell chromatin and transcriptomic profiling have revealed early epigenomic alterations in at-risk individuals relative to RA patients and healthy controls [27, 46]. James et al. reported widespread epigenetic dysregulation in peripheral B cells, memory T cells, and naive T cells from ACPA⁺ at-risk individuals based on DNA methylation profiling [30]. Prideaux et al. further identified baseline methylation loci that distinguished Converters from Nonconverters, enriched in B cell genes related to NOTCH signaling and DNA repair, consistent with the stress-associated phenotype observed in aNAVs in our study [31]. Longitudinal analyses revealed stable methylomes in Nonconverters and ACPA⁻ Controls, mirroring the stable transcriptional profiles seen here. A recent PBMC scRNA-seq study also demonstrated systemic activation signatures in naive B and T cells among ACPA^+^ at-risk individuals compared with healthy controls [32]. However, these prior studies often overlooked the heterogeneity within the ACPA⁺ at-risk population. Here, we demonstrate that ACPA⁺ asymptomatic individuals comprise at least two distinct subgroups: those who progress to RA (Converters) and those who remain RA-free (Nonconverters). Our findings identify a pathogenic naive B cell subset—senescent, polyreactive aNAV^CXCR5⁺^ cells—that distinguishes Converters and Nonconverters at the pre-clinical RA phase. This subset emerges before clinical onset, remains developmental arrested during disease progression, and may functionally contribute to RA conversion.

Recent clinical trials of anti-CD19 CAR-T cell therapies in autoimmune diseases have demonstrated remarkable efficacy, reinforcing the central role of B cells in autoimmune pathogenesis [6, 87, 88]. However, while these therapies show that in-depth B-cell depletion can induce a stable drug-free remission, they do not elucidate how B cells initiate or sustain autoimmune activation, particularly during the pre-clinical phase. Notably, new local tissue toxicity syndromes have been reported in CAR-T–treated patients, likely due to indiscriminate depletion of B cells within affected organs [89]. This underscores the urgent need to identify and selectively target the specific pathogenic B cell subsets that drive disease onset and progression. Previous studies have primarily focused on antibody-secreting B cells as the main pathogenic drivers [90]. In contrast, our findings suggest that senescent and anergy-like aNAV B cells may also represent a key pathogenic subset contributing to disease initiation. Although aNAVs are not active antibody producers, they may promote systemic autoimmune activation through reshaping the global B cell transcriptome and repertoire, releasing proinflammatory cytokines, engaging in aberrant cell–cell interactions, and presenting autoantigens to T cells. Whether selective depletion of senescent aNAV B cells can ameliorate autoimmune flares or reduce disease onset risk remains an important question for future investigation.

We acknowledge several limitations in this study. First, the cohort of ACPA^+^ at-risk individuals was relatively small, with baseline samples collected on average approximately one year before the onset of clinical RA. This limited our ability to pinpoint when, along the pre-clinical disease continuum, the molecular and cellular changes identified here first arise. Second, we only evaluated ACPA^+^ at-risk individuals who were female and post-menopausal; as such, these findings need to be explored in populations that include broader age ranges, and sexes and menopausal states. Third, our B cell isolation strategy failed to capture peripheral plasmablasts. This absence of plasmablasts has also been reported in other studies using the same commercial isolation kit [91], likely due to the kit’s limited capacity to enrich plasmablasts or the inherent fragility of plasmablasts during PBMC isolation, transport and processing. Whether plasmablasts, particularly those clonally related to aNAVs, contribute to RA development at the preclinical stage remains to be determined. Fourth, due to fundamental differences between human and mouse immune systems [92], there is currently no transgenic mouse model that fully recapitulates the ACPA⁺ at-risk phase or the pathogenic expansion of aNAVs prior to RA onset. Whether overexpression or knock-in of key regulators such as KLF2 can drive aNAV accumulation upon antigen stimulation warrants future investigation in both mouse models and human in vitro systems. Fifth, in our in vitro antigen-presentation assays, B cells were pre-stimulated with a cocktail of citrullinated autoantigens, which may have induced broader T cell activation than would be expected from a single defined epitope. This likely represents a polyclonal recall response, reflecting prior antigen exposure in ACPA⁺ individuals. Whether individual citrullinated antigens differ in their ability to activate aNAVs and drive downstream T cell activation in an antigen-specific manner remains to be elucidated. Finally, our analyses were restricted to peripheral B cells. It remains unclear whether intrinsic molecular alterations begin earlier in B cell development within the bone marrow, potentially contributing to the persistent release of polyreactive autoreactive naive B cells. Future studies are needed to address this question.

In summary, this study identifies a B cell subset that is selectively expanded in ACPA⁺ Converters, but not in ACPA+ Nonconverters or ACPA-Controls. These cells display a proinflammatory, senescent phenotype, produce polyreactive autoreactive IgM, and serve as APCs that present citrullinated antigens to activate cytotoxic T cells that together mediate synovitis and synovial tissue destruction in RA. Together, these findings offer mechanistic insight into how naive B cell activation and senescence contribute to the clinical onset of autoimmune disease. They also highlight aNAVs as potential targets for diagnostic and therapeutic strategies aimed at the early prediction and prevention of RA.

## Method and Materials

### TIP-RA cohort

The Targeting Immune Responses for Prevention of Rheumatoid Arthritis (TIP-RA) cohort was designed to prospectively study individuals at high risk for developing rheumatoid arthritis (i.e., At-Risk) due to the presence at baseline of serum anti-citrullinated protein antibody (ACPA) positivity in the absence of a history or physical examination evidence of inflammatory arthritis (IA) at their baseline visit. To identify ACPA^+^ individuals for this study, we used the anti-cyclic citrullinated peptide-3 (anti-CCP3) assay (Inova Diagnostics, Inc., San Diego, California, USA), and the manufacturer’s suggested cut-off level of >=20 units to determine positivity. At-Risk individuals as well as anti-CCP3(−) healthy controls (Controls) were recruited through screening of health-fair participants, first-degree relatives of patients with RA, and individuals referred for evaluation in rheumatology clinics. Of note, anti-CCP3 positive subjects were only included if they tested positive for anti-CCP3 on two occasions: at screening and at their baseline study visit; in addition, anti-CCP3(−) subjects were included if they tested negative for anti-CCP3 on two occasions: screening and baseline study visit.

### Clinical phenotyping (TIP-RA cohort)

All participants in the TIP-RA cohort were evaluated at two study sites: the University of Colorado Anschutz Medical Campus, Aurora, Colorado, USA and the Benaroya Research Institute, Seattle, Washington USA, with recruitment from 2016-2018. All data included in these analyses were obtained at a baseline study visit, and a follow-up visit for all participants. Clinical data was obtained using questionnaires that have been established in previous studies and included self-report of sex assigned at birth. All participants underwent a 66/68 joint examination for tender and swollen joints by a trained examiner. In addition, the absence of IA at baseline in the At-Risk and anti-CCP3(−) participants was confirmed on study-based physical examination as well as study questionnaires and review of their medical records. All study participants were followed longitudinally at an annual visit, or an ‘unscheduled’ visit if they developed new symptoms that were suggestive of development of IA. The outcome of clinical RA was based on clinical evaluation and a finding on physical examination by a rheumatologist or trained study nurse of ≥1 swollen joint consistent with IA.

### Clinical autoantibody and biomarker testing (TIP-RA cohort)

As mentioned above, the primary inclusion ACPA autoantibody biomarker was the anti-CCP3 assay (IgG, Inova Diagnostics Inc., San Diego, California, USA) with positivity determined by the manufacturer’s suggested cut-off level of >=20 units. Additional autoantibody testing including rheumatoid factor (RF) IgA and IgM (Inova Diagnostics Inc.) with positivity determined using a local cut-off level that is equivalent to a level present in <2% of a non-RA population. All testing for these autoantibodies for all subjects was performed in the Exsera Biolabs at the University of Colorado Anschutz Medical Campus (www.exserabiolabs.org). High sensitivity C-reactive protein (hsCRP) testing was performed using nephelometry with results in milligrams per liter (mg/L).

### Final participants (TIP-RA cohort)

For this study, we evaluated a subset of ten participants who exhibited ACPA positivity by the commercial anti-CCP3 ELISA assay (IgG Inova Diagnostics, Inc., San Diego, CA; positive level >=20 units) and who at their baseline study visit did not have historical or examination evidence of IA on a 66/68 joint count by a trained rheumatologist or study nurse. Furthermore, for these ten anti-CCP3+ individuals, we selected five individuals who were known to develop clinically apparent IA (i.e. clinical RA) during the study (Converters) that was further classified as RA by the 2010 American College of Rheumatology/European Alliance of Associations for Rheumatology (ACR/EULAR) Classification Criteria [34]. To minimize variability between participants in the time span of the temporal development of clinical RA, we selected a baseline sample for Converters that was approximately one-year prior to the onset of clinical RA, and a sample from the visit at which clinical RA was first identified. We also selected five anti-CCP3+ individuals (Nonconverters) who did not develop IA/clinical RA during the study; furthermore, to avoid inadvertently including samples from Nonconverters who may have developed clinical RA shortly after their second sample, we selected samples from Nonconverters who were known to have not developed clinical RA for at least 1 year after the second sample was collected. In addition, 3 anti-CCP3(−) individuals without IA or anti-CCP3 positivity were evaluated; these participants are designed anti-CCP3(−) controls. We matched Converters, Nonconverters and Controls by age, race, tobacco use (Supplementary Table 1); in addition, to minimize effects of sex and menopausal status on biologic findings, we only evaluated participants who self-reported female sex at birth, and who self-reported being post-menopausal at the time of their baseline evaluation.

### Ethical considerations (TIP-RA cohort)

The study was reviewed and approved by institutional review boards at all participating institutions including the University of Colorado, Denver, Colorado USA and Benaroya Research Institute, Seattle, Washington USA where participants were evaluated. In addition, the study protocols were approved at Stanford University, Palo Alto, California USA. All participants completed a written informed consent process prior to study participation. All participants received compensation for their participation in study visits.

### PBMC isolation and B cell enrichment (TIP-RA cohort)

Blood from each study participant was drawn into CPT tubes (BD362753 glass with Ficoll Hypaque Solution, draw volume 8 mL) and inverted at room temperature until processing which was completed within 2 hours of initial collection. CPT tubes were centrifuged at 1500-1600G for 30 minutes at room temperature. After centrifugation, plasma was aspirated from the top of the tube and the PBMC layer from up to three CPT tubes was transferred to a 50 mL conical tube (no more than three CPT tubes are pooled in a single 50 mL conical tube). Each conical tube was filled to 50 mL with PBS without calcium2+ and magnesium2+ (Corning 21-031-CM). The conical tube(s) were then centrifuged at room temperature and 800G for 7 minutes. After this second centrifugation, the plasma/PBS solution was aliquoted to 1 cm above the cellular layer, and the cells were resuspended by adding 1 mL of PBS without calcium2+ and magnesium2+. This mixture of cells and PBS was then placed into a new 50 mL conical tube that was filled to 50 mL with PBS (same as above), cells were counted, and the tube was centrifuged at room temperature at 300G for 5 minutes. Excess plasma/PBS was aspirated to 1 cm above the cellular layer, and cells were resuspended in Recovery Cell Culture Freezing Medium in the same 50 mL conical tube. PBMCs were then aliquoted to cryovials with an estimated 10 million cells per cryovial; then, in each cryovial and additional 1 mL of Recovery Cell Culture Freezing Medium was added. Cryovials were then placed into a Mr. Frosty (Sigma C1562) (with isopropanol per manufacturer’s specifications) that was prechilled for at least 8 hours to 4 degree C and then incubated in a 4 degree C refrigerator for 15 minutes. The Mr. Frosty was then transferred to a minus 80 degree C freezer for a minimum of 12 hours (but no longer than 72 hours) and then the cryovials were transferred to liquid nitrogen for long-term storage.

Upon thawing, dead cells were removed using the EasySep™ Dead Cell Removal (Annexin V) Kit (StemCell Technologies, Cat#17899) following the manufacturer’s instructions. Viable cells were then subjected to magnetic negative selection to enrich for B cells using the EasySep™ Human B Cell Isolation Kit (StemCell Technologies, Cat#100-0971). In this procedure, unwanted non-B cells expressing CD2, CD3, CD14, CD16, CD36, CD43, CD56, CD66b, and GlyA were labeled with antibody complexes and magnetic particles. These labeled cells were removed, resulting in a highly purified population of untouched B cells for downstream applications.

### Single-cell multi-omics profiling

Single-cell multi-omics analysis was performed using the 10x Genomics Chromium Single Cell 5’ Reagent Kit v2 (Dual Index), following the manufacturer’s protocol. Briefly, enriched B cells were loaded onto the Chromium X instrument to generate gel bead-in-emulsions (GEMs) for partitioning and barcoding of single cells. Each sample underwent simultaneous 5’ scRNA-seq, cell surface protein profiling via CITE-seq, and V(D)J repertoire sequencing. For CITE-seq, cells were stained with a panel of oligo-conjugated BioLegend TotalSeq™-C antibodies targeting canonical B cell surface markers (listed in Supplementary Table 9) prior to GEM generation. After GEM recovery and reverse transcription, cDNA and antibody-derived tag (ADT) libraries were amplified and processed according to the 10x Genomics protocol. Separate libraries were generated for gene expression (GEX), ADTs, and V(D)J regions. Library quality and concentration were assessed using the Agilent 4200 TapeStation system (Agilent Technologies). Sequencing was performed on the Illumina NovaSeq X Plus platform (Novogene) and the data were processed using Cell Ranger (v9.0.0) and aligned to the Human GRCh38 reference genome.

### Computational analysis of scRNA-seq datasets

scRNA-seq data were analyzed in R using Bioconductor packages [93, 94] and custom scripts. Raw count matrices from individual samples were imported and combined into a unified SingleCellExperiment object using DropletUtils [95] and scMerge [96]. Quality control was performed using a combination of methods. Cells were filtered based on a median absolute deviation (MAD) threshold using the isOutlier function from Scuttle [97]. Additional filtering removed cells with more than 10% mitochondrial gene content, total UMI counts less than 1,000 or greater than 20,000, and total detected genes fewer than 800 or more than 7,000. After filtering, the UMI count matrix was normalized and log-transformed using a deconvolution-based normalization strategy implemented in the scran package [98]. Dimensionality reduction was carried out by performing principal component analysis (PCA) on the top 2,000 highly variable genes. The first 50 principal components were used to generate Uniform Manifold Approximation and Projection (UMAP) embeddings for visualization. Batch correction and sample integration were achieved using the fast mutual nearest neighbor (FastMNN) method from the batchelor package [99]. Cell clustering was performed using a k-nearest neighbor graph-based approach with a range of k values (from 10 to 50), followed by community detection using the Louvain algorithm [100]. B cell clusters were annotated based on the expression of canonical gene markers and surface protein markers derived from CITE-seq data.

### Gene and pathway analyses

For differential gene expression (DEG) analysis, pseudo-bulk expression profiles were generated by summing raw single-cell counts per sample using the Scuttle package [97]. DEG testing was then conducted using the edgeR framework [101], employing Trimmed Mean of M-values (TMM) normalization and likelihood ratio testing (LRT). To interpret the functional relevance of DEGs, pathway enrichment analysis was performed using the clusterProfiler package [102], referencing both GO and KEGG pathway databases. In addition, UCell [103] was used to calculate gene module scores, including senescence score, activation score, and IL-17 pathway score, based on curated gene sets listed in Supplementary Tables.

### BCR repertoire analysis

BCR contigs were initially annotated using the Cell Ranger VDJ pipeline (10x Genomics) and subsequently re-annotated against the IMGT human VDJ reference database using the Change-O package [104]. BCR clonotypes were defined based on paired heavy and light chain sequences with identical V and J gene usage, CDR3 length, and at least 75% similarity in CDR3 amino acid sequence. Clonal expansion was quantified for each sample by calculating the absolute count and relative proportion of each clonotype among total BCR clonotypes. Processed BCR repertoire data were analyzed in R using immunarch, Dowser [105], and custom scripts. A V–J usage matrix was constructed, capturing the number of cells per V–J rearrangement per patient sample.

To characterize clonal diversity, we computed the Shannon diversity index and the number of unique clones using the vegan package [106]. PCA was then performed on the V–J feature matrix to visualize repertoire differences across RA disease stages and clinical conditions. For clonal evolution analysis, BCR clonotype phylogenetic trees were inferred using igphyml [107]. Germline trees connecting distinct clonotypes within individual samples were generated through multiple sequence alignment with MUSCLE [100], followed by tree estimation using RAxML [108]. Final BCR repertoire trees were constructed by integrating clonotype and germline phylogenies, and visualized using the ggtree package [109].

### Selection and expression of recombinant mAbs

Variable heavy and light chain sequences were selected from IgM⁺ aNAV^CXCR5⁺^ B cells with high activation scores, representing samples from healthy controls, non-converters, and converters. For recombinant expression, the selected variable regions were codon-optimized, synthesized (GenScript), and cloned into custom in-house expression vectors containing the human IgG1 constant region for the heavy chain and either the κ or λ constant region for the light chain. Recombinant antibodies were expressed using the TurboCHO™ High-Performance Expression Platform (GenScript) and subjected to quality control by liquid chromatography–mass spectrometry (LC-MS).

### ELISA

Enzyme-linked immunosorbent assays (ELISAs) were performed using Pierce™ 96-Well Polystyrene Plates (Thermo Fisher, Cat#15041). Plates were coated overnight at 4 °C with 1 μg/mL of antigen in carbonate–bicarbonate buffer (Thermo Fisher, Cat#CB01100). The antigen panel included EBNA1 (Abcam, Cat#ab138345), insulin (Thermo Fisher, Cat#RP-10908), LPS (MCE, Cat#HY-D1056), native collagen type II (Cayman, Cat#39125), citrullinated collagen type II (Cayman, Cat#39126), native glucose-6-phosphate isomerase (G6P; Cayman, Cat#18279), citrullinated G6P (Cayman, Cat#30967), native vimentin (Cayman, Cat#11234), citrullinated vimentin (Cayman, Cat#21942), native fibrinogen (Invitrogen, Cat#RP-43142), citrullinated fibrinogen (Cayman, Cat#400076), native histone H2B (Cayman, Cat#11081), and citrullinated H2B (Cayman, Cat#30133). Native and in vitro–citrullinated lysates of Prevotella species (V. pavula) were prepared according to our previously published protocol. After antigen coating, plates were washed six times with PBST (PBS containing 0.05% Tween-20) and blocked with SuperBlock™ Blocking Buffer (Thermo Fisher, Cat#37515) for 1 h at room temperature. Recombinant monoclonal antibodies (mAbs) were diluted in blocking buffer to a final concentration of 10 μg/mL, added to each well, and incubated for 2 h at room temperature. Plates were then washed six times with PBST and incubated for 1 h at room temperature with HRP-conjugated goat anti-human IgG (H+L) secondary antibody (Thermo Fisher, Cat#62-8420; 1:4000 dilution in blocking buffer). Following six additional PBST washes, TMB substrate (Thermo Fisher, Cat#34028) was added, and the reaction was developed for 15 min at room temperature in the dark. For titration analyses, the incubation time was shortened to 10 minutes to distinguish binding at high versus low antibody concentrations. Reactions were stopped with Stop Solution (Thermo Fisher, Cat#N600), and absorbance at 450 nm (OD450) was measured using a BioTek Synergy H1 microplate reader. Each ELISA plate included a blank control, a non-polyreactive mAb, and a known polyreactive mAb to control for background and normalize inter-plate variability. mAbs were considered antigen-positive if OD450 exceeded 0.7, based on comparison with negative controls. Polyreactive mAbs were defined as antibodies binding to ≥ 2 distinct antigens above the OD450 threshold. OD450 values were visualized as heatmaps using GraphPad Prism (version 10).

### Synovium tissue collection and processing (Stanford and HSS cohort)

Cryopreserved synovial tissue samples from osteoarthritis (OA) or ACPA⁺ rheumatoid arthritis (RA) patients were obtained from the VA Palo Alto Health Care System (Stanford IRB 3780) or the Hospital for Special Surgery (HSS IRB 2014-233) (Supplementary Table 6). At the time of sampling, all RA patients had active disease, defined by a Disease Activity Score (DAS) > 2.6. For tissue dissociation, samples were thawed completely at 37 °C and washed in RPMI 1640 medium (Corning Life Sciences) supplemented with 10% fetal bovine serum (FBS; ATCC) and 1% L-glutamine (Gibco). Tissues were minced and digested in RPMI 1640 containing 100 µg/mL Liberase TL (Roche) and 100 µg/mL DNase I (Roche) at 37 °C for 30 min, with intermittent inversion to facilitate dissociation into single-cell suspensions. The digested material was passed through a 70 µm cell strainer and washed with cold RPMI 1640 containing 10% FBS and 1% L-glutamine. B cells were further enriched using the EasySep™ Human B Cell Isolation Kit (StemCell Technologies, Cat#100-0971). Single-cell suspensions were counted, prepared for antibody staining, and processed for 10x Genomics single-cell RNA sequencing. Libraries were generated on a Chromium Controller (10x Genomics) using the Chromium Next GEM Single Cell 5’ v2 (Dual Index) kit with Feature Barcode technology, enabling both gene expression and V(D)J repertoire profiling, following the manufacturer’s instructions.

### ACPA^+^ RA PBMCs (Stanford and VA Palo Alto cohort)

All ACPA⁺ RA samples used for in vitro studies were obtained under institutional review board–approved protocols (IRB #3780) at the Veterans Affairs Palo Alto Health Care System and Stanford University (Supplementary Table 8). Written informed consent was obtained from all participants prior to blood collection. Healthy donor samples were randomly acquired from the Stanford Blood Center. Whole blood was collected in heparinized Vacutainer tubes (BD Biosciences) and processed to isolate PBMCs by Ficoll-Paque density gradient centrifugation (Sigma-Aldrich) using Leucosep™ tubes (Greiner Bio-One). Isolated PBMCs were cryopreserved in Recovery™ Cell Culture Freezing Medium (Thermo Fisher Scientific) until use.

### Flow cytometry and cell sorting

Following cell culture or PBMC thawing, cells were pelleted at 350 × g for 5 min and washed with PBS to obtain single-cell suspensions. Cells were resuspended in FACS Stain Buffer (BD Biosciences) supplemented with Fixable Viability Stain 510 (BD Biosciences) and Human TruStain FcX (BioLegend), and incubated for 15 min at 4 °C. After washing with stain buffer, cells were labeled for 30 min at 4 °C with fluorescently conjugated antibodies targeting surface markers. For intracellular staining, cells at the end of isolation or culture were treated with eBioscience Protein Transport Inhibitor Cocktail (Thermo Fisher Scientific) during the final 5 h of culture. Cells were then washed, fixed, and permeabilized using the eBioscience Fixation/Permeabilization Kit (Thermo Fisher Scientific), followed by a 30 min incubation at 4 °C with antibodies against intracellular targets. Flow cytometry was performed on a BD LSRFortessa (BD Biosciences), and data were analyzed using FlowJo software (BD Biosciences).

For B cell subset and CD3⁺ T cell sorting from ACPA⁺ RA donors, cryopreserved PBMCs were thawed at 37 °C and washed in FACS Stain Buffer to generate single-cell suspensions. Cells were stained with Fixable Viability Stain 510 and TruStain FcX for 15 min, followed by labeling with anti-CD19, IgD, CD27, IgM, CXCR5, CD69, and CD3 antibodies in FACS Stain Buffer. Sorting was performed using a BD FACSAria II (BD Biosciences) or Sony SH800 (Sony Biotechnology) cell sorter. Antibodies used for staining are listed in Supplementary Table 10.

### Coculture experiments

Bulk resting naive (rNAV), activated naive (aNAV), or CD27⁺ CD19⁺ B cells, along with CD3⁺ T cells, were isolated from ACPA⁺ RA PBMCs using the cell sorting strategy described above. B cells were pre-stimulated for 3 h with 5 μg/mL anti-IgM antibody (Jackson ImmunoResearch) and 5 μg/mL native or citrullinated proteins—including histone H3, vimentin, and fibrinogen (Cayman Chemical)—in the presence or absence of 5 μg/mL anti-HLA class I/II antibodies (BioLegend) to block HLA-mediated antigen presentation. Following stimulation, B cells were cocultured with T cells at a 1:10 ratio for 7 days in Iscove’s Modified Dulbecco’s Medium (IMDM; Thermo Fisher) supplemented with 10% FBS, 1% penicillin–streptomycin, 1% L-glutamine, 2 μg/mL anti-CD28 antibody (BD Biosciences), 10 ng/mL IL-2 (Thermo Fisher), 50 ng/mL IL-21 (Thermo Fisher), and 100 ng/mL recombinant BAFF (R&D Systems). After 7 days, T and B cell proliferation and activation were assessed by flow cytometry.

### B cell activation and senescence assays

For the naive B cell activation assay, CD3^-^CD19^+^CD27^-^IgM^+^IgD^+^CXCR5^-^CD69^-^ rNAV B cells were isolated from ACPA⁺ RA PBMCs by flow cytometric sorting and cultured in complete IMDM medium supplemented with 10% FBS, 1% penicillin–streptomycin, and 1% L-glutamine. Cells were stimulated for 2 days with 5 μg/mL anti-IgM antibody (Jackson ImmunoResearch), 5 μg/mL native or citrullinated proteins (Cayman Chemical), 1 μg/mL LPS (Sigma-Aldrich), or 1 μg/mL CpG oligodeoxynucleotides (InvivoGen). Activation was assessed by quantifying CXCR5, CD69, and CD86 expression within the CD19⁺IgM⁺IgD⁺CD27⁻ population by flow cytometry. For the senescence assay, sorted B cells were stimulated for 7 days with the same antigenic conditions described above. Cells were stained with antibodies against CD19, CD27, IgD, IgM, CD38, CD69, and CXCR5 for 30 min at 4°C, fixed with 2% paraformaldehyde for 10 min at room temperature, and washed with FACS buffer. Senescence levels were determined using the CellEvent™ Senescence Green Flow Cytometry Assay Kit (Thermo Fisher, Cat#C10841) according to the manufacturer’s instructions, with a 2 h incubation at 37°C prior to flow cytometric analysis.

## Data analysis and statistics

Statistical analyses were performed using GraphPad Prism (version 10), R (version 4.3.2), and Python (version 3.10). Data in bar graphs and dot plots are presented as mean ± standard deviation (s.d.), unless otherwise noted. Individual data points represent independent biological replicates (e.g., patient samples or single cells). Data distribution was assessed using the Kolmogorov–Smirnov (K–S) test. For data that followed a normal distribution, comparisons between two groups were performed using two-tailed unpaired Student’s t-tests, and comparisons among multiple groups were conducted using one-way ANOVA followed by Tukey’s post hoc test. For normally distributed data involving two independent variables (e.g., time and condition), two-way ANOVA was applied. For non-normally distributed data, the Mann–Whitney U test (for two groups) or Kruskal–Wallis test followed by Dunn’s post hoc test (for multiple groups) was used. Gene Ontology (GO) and KEGG pathway enrichment analyses were conducted using the clusterProfiler package (version 4.6.2). P-values were adjusted using the Benjamini–Hochberg method to control the false discovery rate (FDR), with adjusted P < 0.05 considered statistically significant. Statistical significance is indicated as follows: P < 0.05 (*), P < 0.01 (**), P < 0.001 (***), and P < 0.0001 (****); ns indicates not significant.

## Data availability

The single cell RNA-Seq datasets generated and analyzed in this study have been uploaded to the Sequence Read Archive (https://www.ncbi.nlm.nih.gov/sra) with the accession number PRJNA1344838 (peripheral B cell sequencing) and PRJNA1345084 (synovial B cell sequencing). Source data are provided with this paper.

## Acknowledgements

This work was supported by the National Institutes of Health (NIH) through the following grants: R01 AR078268, R01 AI173189-01, and PATHO-PH2-SUB_17_23 (all to W.H.R.).

Additional support was provided by the Brennan Family, the Arthritis Research Coalition (ARC) (to W.H.R.), and the Stanford Dean’s Postdoctoral Fellowship (to X.W.). The research reported in this publication was also supported by NIH/NIAMS P30 AR079369 (to V.M.H., G.S.F., K.D.D., and L.M.); the William P. Arend Endowed Chair for Rheumatology Research (to K.D.D.); and the Autoimmune Disease Prevention Center at the University of Colorado Anschutz Medical Campus (to G.S.F., K.D.D., and L.M.). This work was further supported by an investigator-initiated research grant from Janssen Research & Development, LLC (to W.H.R., V.M.H., K.D.D., L.M., and Y.O.).

## Author contributions

X.W. conceived and designed experiments, performed experiments, analyzed data, and wrote the manuscript. M.Z., J.S.M., E.K.S., and J.C.A. performed experiments and analyzed data. O.S., M.F., L.M., M.H.S., L.T.D., J.H.B., E.A.J., G.S.F., Y.O., V.M.H., and K.D.D. collected human samples. L.S.v.D., P.P.H., T.V.L., E.M., V.M.H., and K.D.D. provided expertise and edited the manuscript. K.D.D. and W.H.R. conceived and designed experiments, analyzed data, and wrote the manuscript.

## Supplemental figures

**Figure S1.**
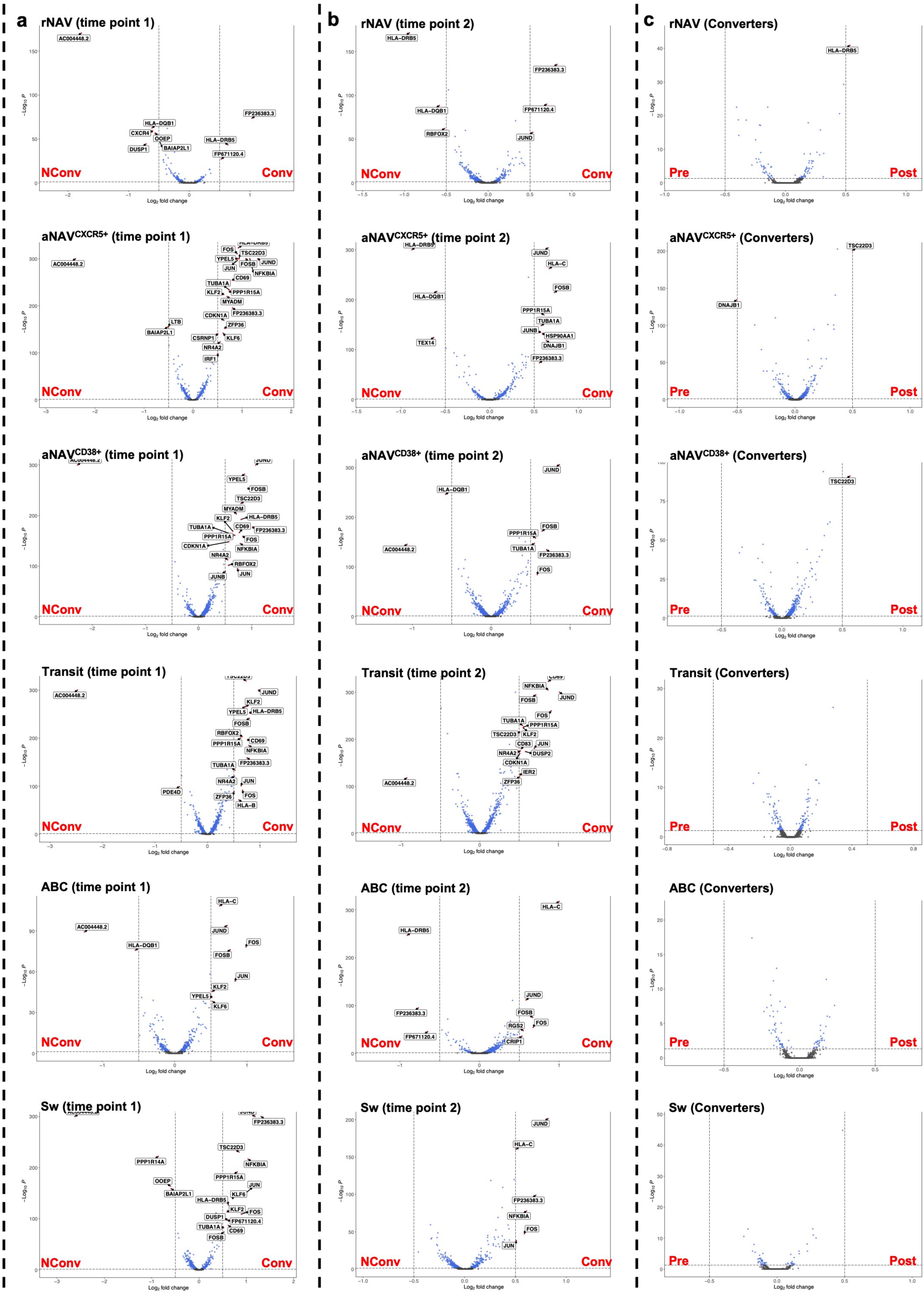
Subgroup differential gene expression (DEG) analysis of the B-cell transcriptome. (a) DEGs comparing Converters and Nonconverters at the pre-clinical time point. (b) DEGs comparing Conv and Nconv at the post-onset time point. (c) DEGs comparing preclinical versus post-onset time points within Converter B-cell subsets. DEGs were identified using the Wilcoxon rank-sum test with Bonferroni correction. Top 20 genes with an average log₂ fold change > 0.5 and adjusted P < 0.05 are labeled in the plots. Conv: Converters; NConv: Nonconverters.

**Figure S2.**
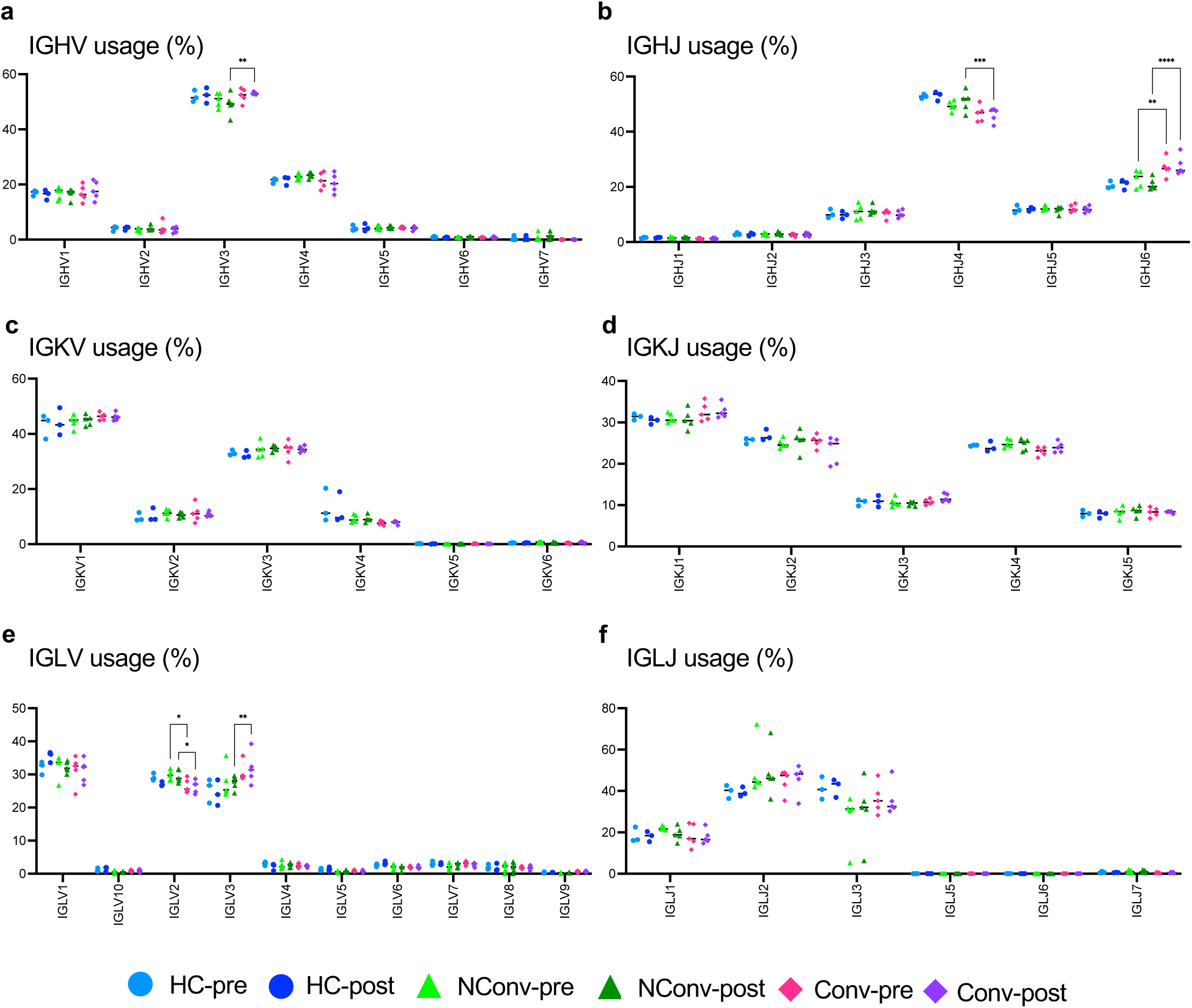
V and J gene usage analysis of BCR heavy and light chains. (a, b) IGHV and IGHJ gene usage in the heavy chains. (c, d) IGKV and IGKJ gene usage in the κ light chains. (e, f) IGLV and IGLJ gene usage in the λ light chains. *P < 0.05; **P < 0.01; ***P < 0.001; ****P < 0.0001 by two-way ANOVA. HC: ACPA^-^ Controls; NConv: ACPA^+^ Nonconverters; Conv: ACPA^+^ Converters.

**Figure S3.**
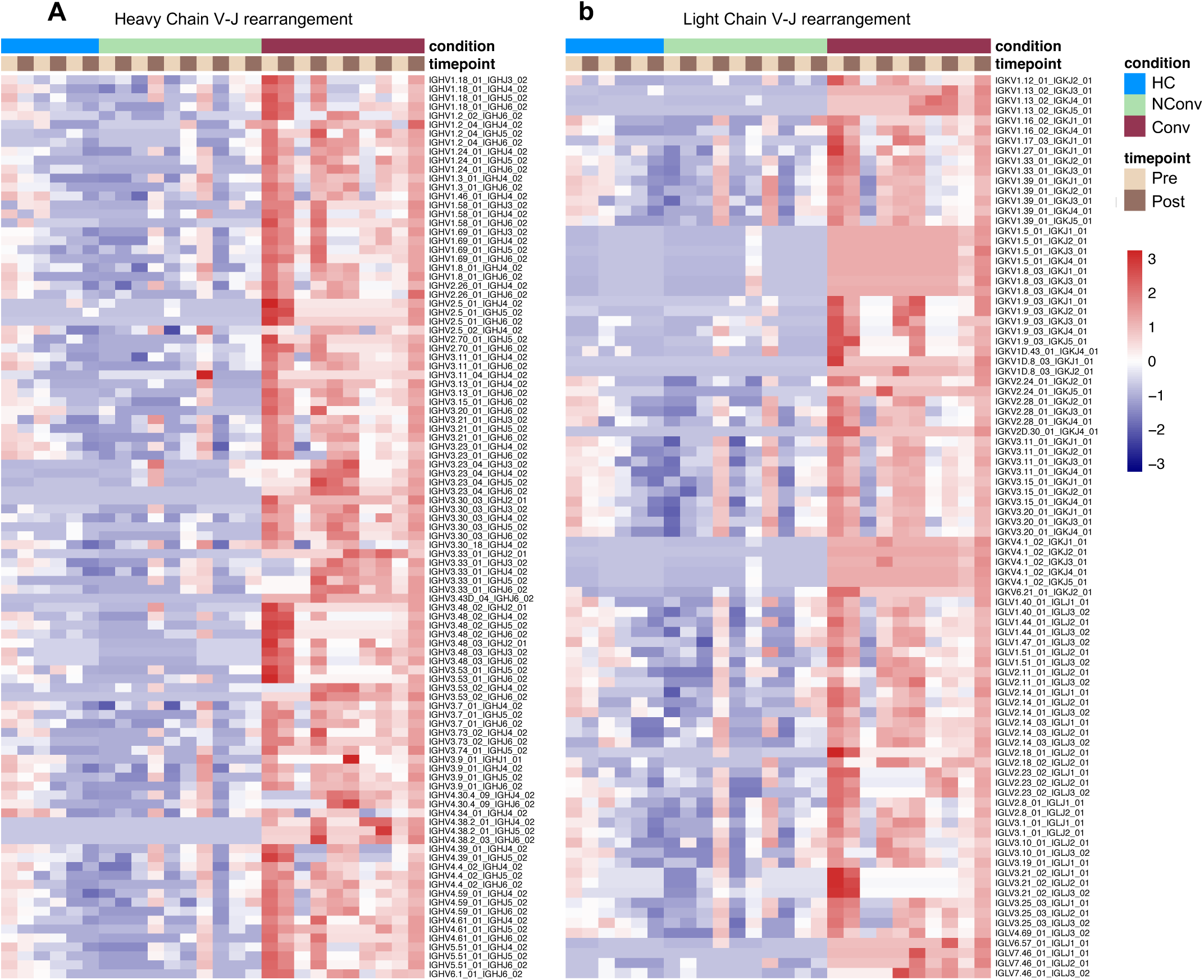
Heatmap of distinct V–J rearrangement patterns in Converters’ aNAV^CXCR5+^ cells. (a, b) Heatmaps showing V–J rearrangement patterns enriched in Converters but not in Nonconverters or Controls. (a) Heavy chain. (b) Light chain. Each column represents a sample of aNAV^CXCR5⁺^ cells from an individual at a given time point, and each row represents a specific V–J rearrangement combination. HC: ACPA^-^ Controls; NConv: ACPA^+^ Nonconverters; Conv: ACPA^+^ Converters.

**Figure S4.**
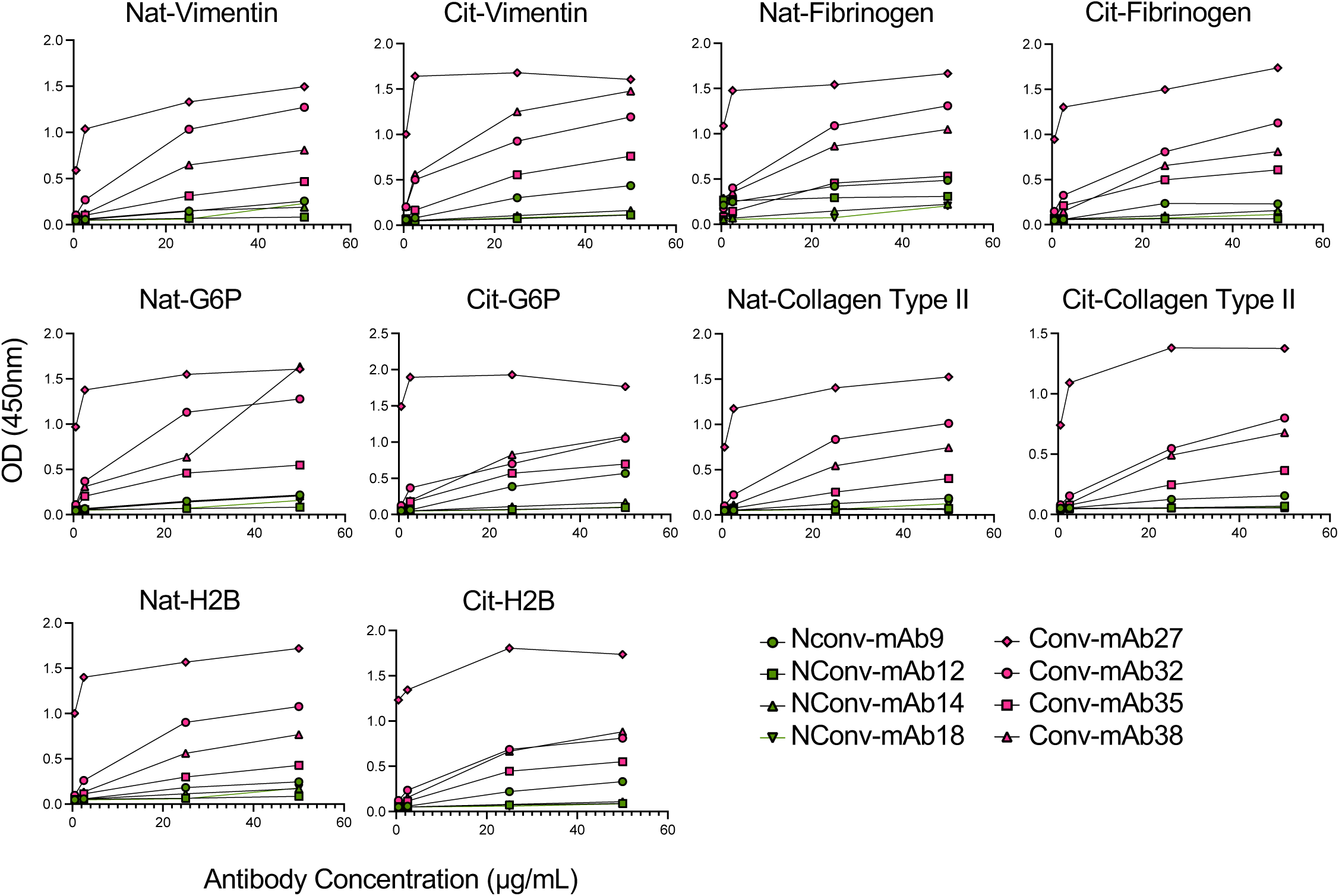
Reactivity analysis of aNAV-derived mAbs to RA antigens in Nonconverters and Converters. Polyreactive mAbs from Converters (mAb27, mAb32, mAb35, and mAb38) and Nonconverters (mAb9, mAb14, and mAb18) were tested against native and citrullinated RA antigens (i.e., vimentin, fibrinogen, G6P, collagen type II, H2B) by ELISA at serial concentrations (2.5, 5, 25, and 50 µg/mL). Nonconverter’s mAb12 was included as a negative control. HC: ACPA^-^ Controls; NConv: ACPA^+^ Nonconverters; Conv: ACPA^+^ Converters.

**Figure S5.**
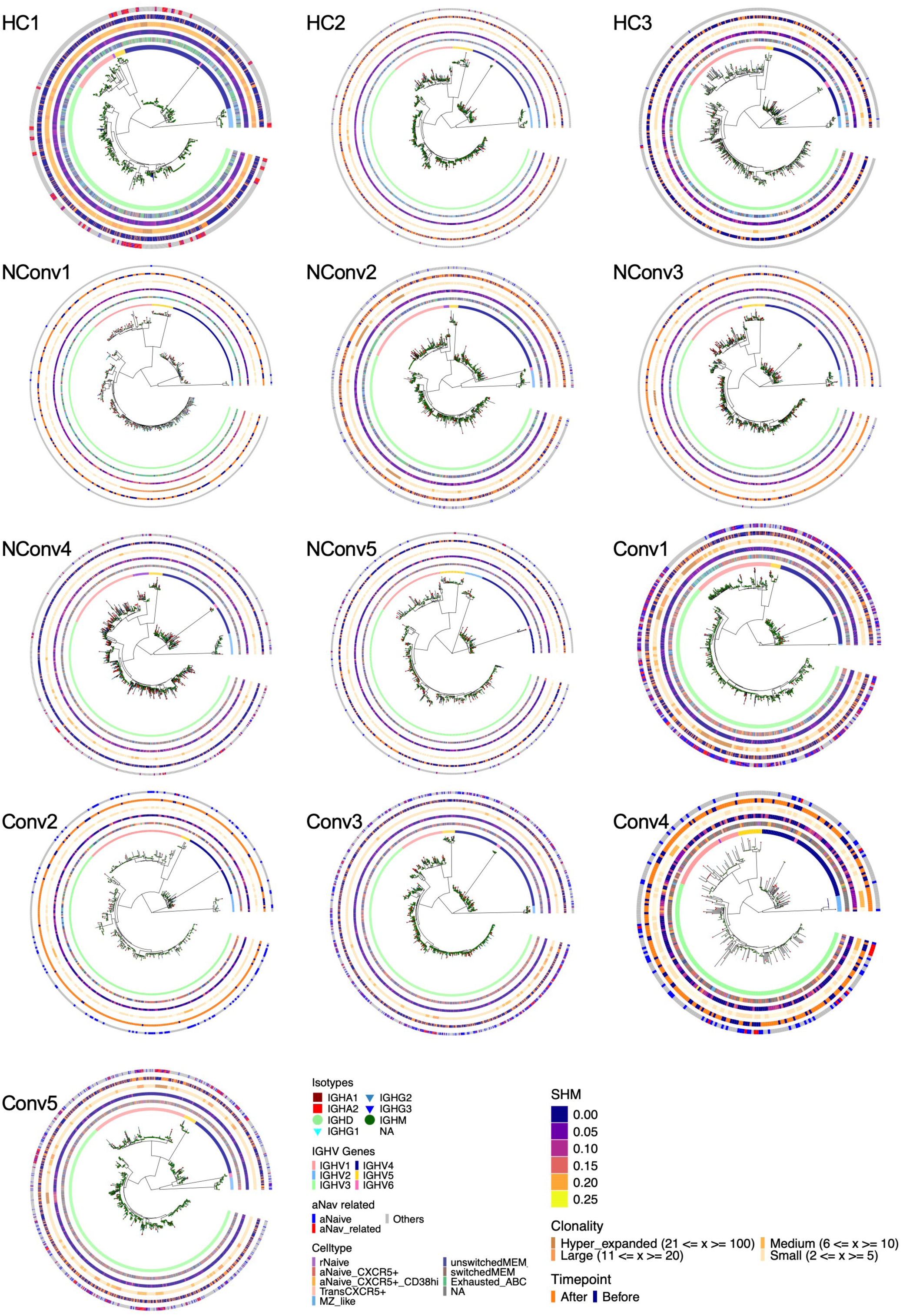
Phylogenetic analysis of the BCR repertoire. Phylogenetic relationships were analyzed based on CDR3 length and sequence similarity. BCR sequences with the same CDR3 length and >75% heavy-chain similarity were defined as clonal families. Information on isotype, V gene, J gene, cell type, and time point is indicated in the plots. HC: ACPA^-^ Controls; NConv: ACPA^+^ Nonconverters; Conv: ACPA^+^ Converters.

